# Electrical coupling controls dimensionality and chaotic firing of inferior olive neurons

**DOI:** 10.1101/542183

**Authors:** Huu Hoang, Eric J. Lang, Yoshito Hirata, Isao T. Tokuda, Kazuyuki Aihara, Keisuke Toyama, Mitsuo Kawato, Nicolas Schweighofer

## Abstract

One of the main challenges facing online neural learning systems with numerous modifiable parameters (or “degrees-of-freedom”) such as the cerebellum, is how to avoid “overfitting”. We previously proposed that the cerebellum controls the degree-of-freedoms during learning by gradually modulating the electric coupling strength between inferior olive neurons. Here, we develop a modeling technique to estimate effective coupling strengths between inferior olive neurons from *in vivo* recordings of Purkinje cell complex spike activity in three different coupling conditions. We show that high coupling strengths induce synchronous firing and decrease the dimensionality of inferior olive firing dynamics. In contrast, intermediate coupling strengths induce chaotic firing and increase the dimensionality of firing dynamics. Our results thus support the hypothesis that effective coupling controls the dimensionality of inferior olive firing, which may allow the olivocerebellar system to learn effectively from a small training sample set despite the low firing frequency of inferior olive neurons.

## INTRODUCTION

Fifty years ago, David Marr proposed a theory that synaptic plasticity in the cerebellar cortex induces motor learning (Marr, 1969). Multiple experimental and computational studies have since supported this theory: climbing fiber afferents carry error signals (Bazzigaluppi et al., 2012; Keating and Thach, 1995; Kitazawa et al., 1998; Kobayashi et al., 1998), that modify parallel-fiber-Purkinje-cell synapses (D’Angelo et al., 2016; Hansel et al., 2001; Ito, 2001; Kuroda et al., 2001), and drive learning of internal models for motor control (Bastian, 2006; Kawato and Gomi, 1992; Schweighofer et al., 1998; Tseng et al., 2007; Vinueza Veloz et al., 2015; Herzfeld et al. 2018). A fundamental question remains unanswered, however: how does the cerebellum learn to control the high dimensional and nonlinear motor systems that are typical of vertebrates for complicated movement patterns, while even the most advanced robots fail to perform similar movements (Adolph et al., 2012; Atkeson et al., 2018)? Moreover, how does the cerebellum achieve such learning despite being constrained by the relatively small numbers of training samples and the low-firing frequency from the inferior olive (IO) neurons, which give rise to the climbing fiber inputs? Here, we address these questions at three distinct levels: *computational*, *algorithmic*, and *implementational* (Marr, 1982), and provide computational and experimental clues for possible answers.

At the computational level, artificial learning systems that have many adjustable parameters require a proportionally large number of training samples to achieve adequate learning generalization (LeCun et al., 2015; Mnih et al., 2015; Silver et al. 2016). In contrast, if the number of training samples is lower than the number of parameters, severe overfitting to the noise in the data occurs. This creates a large generalization error, proportional to *D*/(2*n*), where *n* is the number of training samples and *D* is the number of degree-of-freedoms (DOFs), which is the number of adjustable parameters in the learning system (Watanabe, 2009; Yamazaki, 2014). Therefore, an online learning system, such as the cerebellum, must keep the number of DOFs small in the early stages of learning when the training sample set is small, and then increase it gradually during the course of learning (Schaal et al., 2002; Garrigues and Ghaoui, 2009).

At the algorithmic level, we earlier proposed that the number of cerebellar DOFs is modulated by the degree of synchrony between IO neurons (Kawato et al. 2011; Schweighofer et al. 2013; Tokuda et al. 2017). According to this proposal, early in learning, IO synchrony is high and groups of related neurons in the olivo-cerebellar system behave, in the limit, as a single-neuron chain, thus decreasing the number of DOFs. The resulting synchronous IO error signals would both significantly improve real-time motor control (Lang et al., 2016), and lead to massive changes in efficacies of the parallel-fiber-Purkinje-cell synapses, resulting in fast but crude learning. As learning of the motor act progresses, IO synchrony is decreased, potentially allowing the occurrence of chaotic resonance to enhance information transmission of the error signals (Schweighofer et al., 2004; Tokuda et al., 2010; Masuda and Aihara, 2002; Makarenko and Llinas, 2005; Nobukawa and Nishimura, 2016), which would overcome the constraint of low IO firing rates (Eccles et al., 1966; Llinás and Yarom, 1981). Specifically, chaotic resonance would increase the number of DOFs, and thereby allow more sophisticated learning, resulting in fine tuning of the motor act (Tokuda et al., 2010, 2013).

At the implementational level, we propose that the distinctive features of the IO neurons, the source of the climbing fibers, and the anatomy of the loop formed by the Purkinje cells, deep cerebellar nucleus neurons, and IO neurons, allow modulation of the number of DOFs during learning. The IO neurons form the strongest electrically coupled neuronal network in the adult mammalian brain (De Zeeuw et al., 1995; Condorelli et al., 1998; Belluardo et al., 2000), with electrical synapses driving synchronization when the coupling is strong (Blenkinsop and Lang, 2006 Marshall et al., 2007; Lang 2002). In addition, presynaptic GABAergic terminals control the efficacy of electrical coupling (Llinas et al., 1974; Sotelo et al., 1974; Best and Regehr, 2009; Onizuka et al., 2013; Lefler et al., 2014). These GABAergic afferents largely arise from the deep cerebellar nucleus (de Zeeuw et al., 1989; Nelson and Mugnaini, 1989; Fredette and Mugnaini, 1991), which are part of the anatomical closed-loops formed between corresponding regions of the IO, cerebellar cortex, and deep cerebellar nuclei (Sugihara and Shinoda, 2004, 2007; Apps and Hawkes, 2009; Sugihara et al., 2009). Thus, the Purkinje cells, via this feedback circuit, can regulate the synchrony levels of their corresponding climbing fiber inputs, through the double-inhibition within the feedback circuit (Marshall and Lang, 2009).

Although we have shown in simulations that decreasing IO synchrony via modulation of electrical coupling enhances cerebellar learning (Schweighofer et al., 2004; Tokuda et al., 2010, 2013), there are no experimental supports to the basic assumptions about how electrical coupling, synchrony, chaotic firing, and dimensionality of firing dynamics are linked. Here, we analyze the effect of coupling on the DOFs and on the induction of chaotic resonance by utilizing *in vivo* recordings of complex spikes under three coupling conditions. Specifically, we examine two predictions of our previous hypotheses that 1) increasing the synchrony level, via increased electrical coupling between inferior olive neurons, decreases the dimensionality of the IO firing dynamics and 2) intermediate coupling induces chaotic spiking and maximizes the dimensionality of inferior-olive firing dynamics.

## RESULTS

### Estimation of the effective coupling between IO neurons *in vivo*

To examine the effect of electrical coupling on dimensionality reduction and chaotic dynamics, we first need to estimate the range of effective coupling strengths between IO neurons. Direct quantitative measurement of electrical coupling in the IO has been obtained in slice preparations (Devor and Yarom, 2002; Hoge et al., 2011; Lefler et al., 2014); however, it remains technically impossible to measure *in vivo*. We thus employed an indirect approach to estimate the coupling that involved comparing Purkinje cell complex spike activity recorded simultaneously from arrays of Purkinje cells (Blenkinsop and Lang, 2006; Lang, 2002; Lang et al., 1996) with simulated activity generated by a model of the IO using a Bayesian method, which we previously proposed and validated (Hoang et al., 2015)^1^. Here, we modified this method to further improve the robustness of the coupling estimations via Bayesian model-averaging.

Briefly, in the model, each IO neuron comprises a soma, a main dendrite, and four dendritic spine compartments, with these compartments having distinct ionic conductances. Most notably, the dendritic compartment has a high threshold calcium conductance and a calcium-activated potassium conductance, which are responsible for the after-depolarization and after-hyperpolarization sequence that follows each sodium spike and for the low firing rates of IO neurons (Schweighofer et al., 1999). Each neuron was coupled to its neighboring neurons via electric coupling conductances on the spine compartments, with one inhibitory input conductance per spine. Synaptic noise was added to better account for stochastic process in IO neurons - for review of IO anatomy and function, see (De Zeeuw et al., 1998) and of the model, see (Hoang et al., 2015; Onizuka et al., 2013).

From the model, it is possible to derive the theoretical “effective” electrical coupling conductance *g*_*eff*_ as a function of the axial conductance of the spines *g*_*s*_, the electrical coupling conductance *g*_*c*_, and the GABAergic synaptic conductance *g*_*i*_ (see Katori et al., 2010 and Experimental Procedures for details). Estimates for *g*_*c*_ and *g*_*i*_, were obtained by comparing the model spike activity to the complex spike data sets (*g*_*s*_ was held constant). However, initial simulations showed that the firing frequency of the synaptic noise inputs significantly affected the spiking behavior of the neurons in the model, thus the fit of the firing dynamics of the model to the data and the estimation results. To address this issue, we estimated *g*_*c*_ and *g*_*i*_ for different values of synaptic noise input frequencies via a model-averaging approach (Grueber et al., 2011). Specifically, we first constructed a number of models with different frequencies of synaptic noise inputs, as observed in cerebellar slice data (Najac and Raman, 2015). We then obtained an estimate of *g*_*i*_ and *g*_*c*_ via the Bayesian estimation method from each model (Hoang et al., 2015; see Experimental Procedures for details). The final estimates of *g*_*i*_ and *g*_*c*_ were obtained by averaging these individual model estimates, weighted in proportion to the goodness-of-fit of the models via Bayesian model-averaging (see Figure S2 and Experimental Procedures for details).

Following the procedures outlined above, estimates of *g*_*i*_ and *g*_*c*_ were obtained for three coupling conditions (low, control, high). The low coupling condition was generated by intra-IO injection of the gap junction blocker carbenoxolone (CBX), whereas the presumed high coupling condition was generated by intra-IO injection of the GABA blocker picrotoxin (PIX). Estimation results show that, as expected, the mean *g*_*i*_ and *g*_*c*_ were reduced approximately 20% and 22% under the PIX and CBX from their own control (CON) values, respectively. When the two CON groups were combined, the estimated inhibitory conductance *g*_*i*_ in the CON condition (Figure 1A – 1.11 ± 0.22 mS/cm^2^, *n* = 90 neurons) was significantly decreased in the PIX condition (0.84 ± 0.26 mS/cm^2^*, n* = 46 neurons, PIX-CON: p < 0.0001) and was comparable to that in the CBX condition (1.02 ± 0.13 mS/cm^2^*, n* = 44 neurons, CBX-CON: p = 0.03). Similarly, the estimated gap-junctional conductance *g*_*c*_ in the CBX condition (Figure 1B – 0.87 ± 0.23 mS/cm^2^, CBX-CON: p < 0.0001) was smaller than in the CON condition (1.22 ± 0.28 mS/cm^2^), but there was no significant difference between the PIX (1.29 ± 0.18 mS/cm^2^, PIX-CON: p = 0.23) and CON conditions.

**Figure 1:**
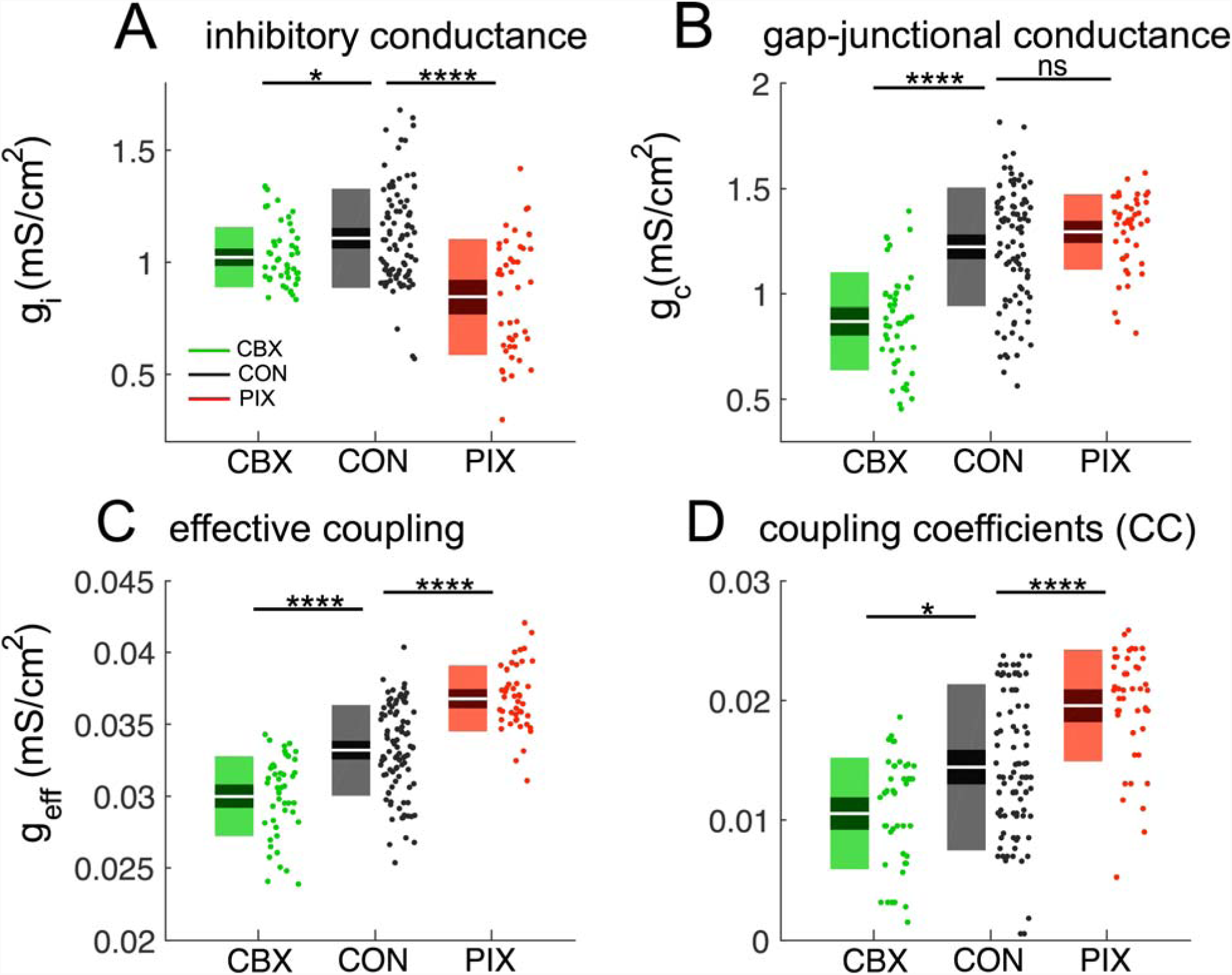
Conductance and coupling coefficient estimation in three experimental conditions. A-B: boxplot with values of *g*_*i*_ (A) and *g*_*c*_ (B) estimates for the three experimental conditions: carbenoxolone (CBX, green box), control (CON, black box) and picrotoxin (PIX, red box). The color conventions are same for subsequent plots. C: the effective coupling *g*_*eff*_ computed as the equation (1) for the three conditions. D: the coupling coefficient (CC) estimated for the three conditions via simulations. Each boxplot shows white line as the mean, dark region as 95% CIs and light region as 1 std. Asterisks represent significance levels: ns p > 0.05, *p < 0.05, ****p < 0.0001.

As a result of these changes in *g*_*i*_ and *g*_*c*_, the estimated effective coupling strength, *g*_*eff*_, calculated using Equation 1, differed across the three conditions (one-way ANOVA: p < 0.0001). It was smallest for the CBX condition (Figure 1C – *g*_*eff*_ = 0.030 ± 0.003 mS/cm^2^, CBX-CON: p < 0.0001), intermediate for the CON condition (*g*_*eff*_ = 0.033 ± 0.003 mS/cm^2^) and largest for the PIX condition (*g*_*eff*_ = 0.037 ± 0.002 mS/cm^2^, PIX-CON: p < 0.0001). The estimated *g*_*i*_ and *g*_*c*_ parameters were then used in the neuronal network to generate simulated spike trains under all three conditions. In each case, the spike trains were comparable to those of the recorded complex-spike activity. In particular, the firing rates of neurons in the CBX condition were lower (see Figure 2A, firing rate of model 0.58 ± 0.55 Hz; data 0.59 ± 0.55 Hz) than in the CON condition (see Figure 2B, model 1.34 ± 0.77 Hz; data 1.36 ± 0.8 Hz), and firing rates in the PIX condition were higher (Figure 2C, model 2.42 ± 0.95 Hz; data 2.51 ± 1.05 Hz).

**Figure 2.**
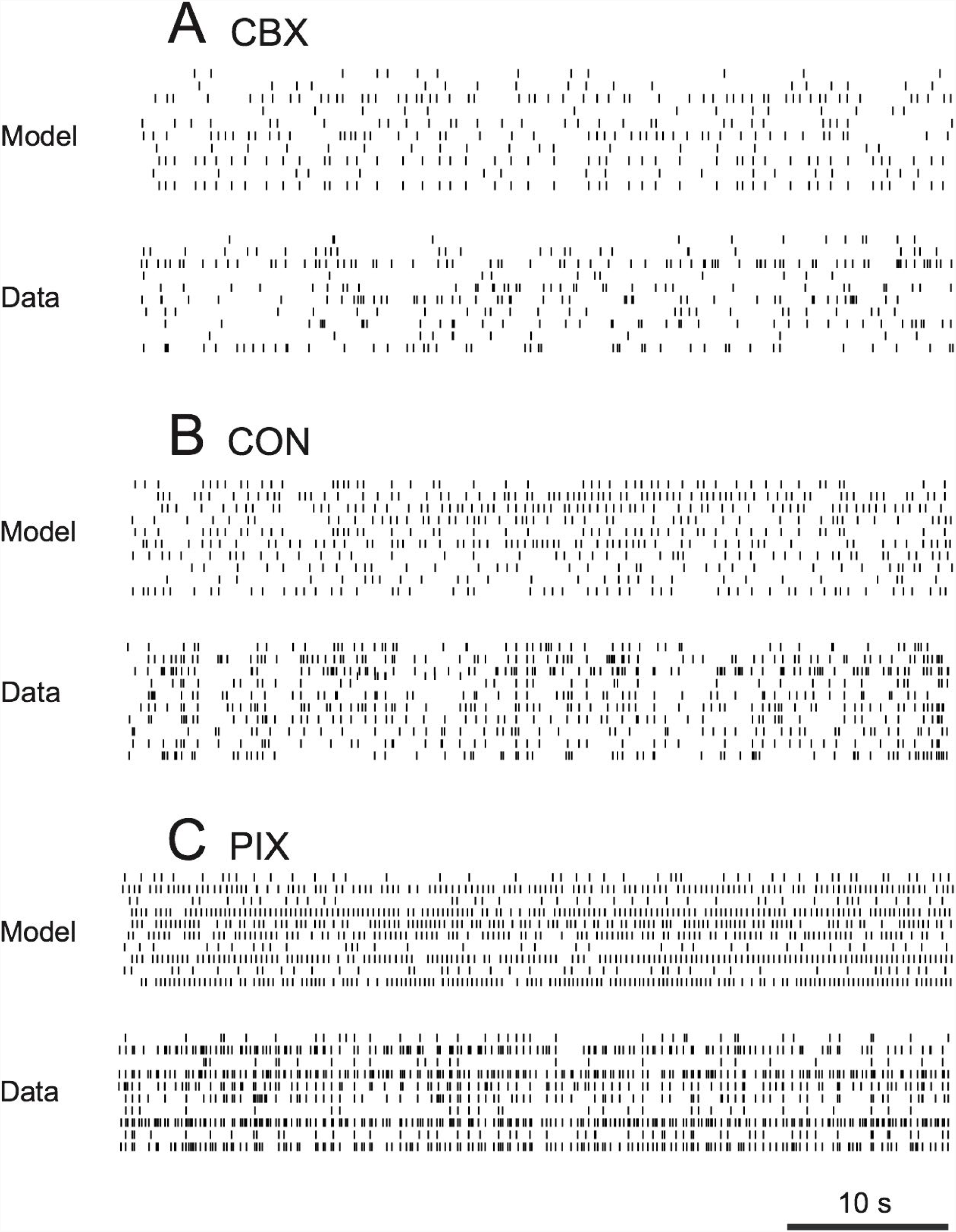
Examples of inferior olive firing for the model and the data. Raster plot of ten representative inferior olive neurons of the model and the experimental complex spike data of three animals in the three conditions. A. Carbenoxolone (animal #1, repetitive spiking). B. Control (animal #7, chaotic) C. Picrotoxin (animal #9, highly synchronous spiking).

Next, to examine whether the estimates for effective coupling strengths in the three conditions were biologically realistic, we computed the coupling coefficients (CCs) for the model neurons. We first hyperpolarized all model neurons to −69 mV by injection of I_hyp_ = −1 μA/ cm^2^ to make them responsive to the stimulus. We then injected a step current I_cmd_ = −1 μA/cm^2^ in the soma of the center neuron and computed the CCs as the average ratio of change in steady state membrane potentials of this “master” cell and its four neighboring cells in the network (Figure S3). As expected, CC was smaller in the CBX condition (Figure 1D – *CC* = 0.01 ± 0.005, CBX-CON: p = 0.01) and larger in the PIX condition (*CC* = 0.02 ± 0.004, PIX-CON: p < 0.0001) than in the CON condition (*CC* = 0.014 ± 0.007). Furthermore, CCs are highly compatible with previously reported *in vitro* values (Lefler et al., 2014). In particular, our results showed that CCs in the PIX and CON conditions were comparable to those of the control (*CC* = 0.021 ± 0.02, cf. Table S1, Lefler et al., 2014) and light-activated (*CC* = 0.012 ± 0.013) conditions, respectively. In both cases, a larger (about double) CC value was found for the condition with less GABA activity (PIX in our experiments and control in Lefler et al, 2014).

### The dimensionality reduction by effective coupling

To quantitatively investigate how synchrony and dimensionality change when effective coupling varies, we estimated the synchrony level and the dimensionality from the spike data of IO neurons in individual animals (see Experimental Procedures for more details). In strong agreement with previous studies (Blenkinsop and Lang 2006; Lang 1996), the synchrony level in 1-ms bins increased 2-3 fold in the PIX condition (synchrony = 0.068 ± 0.051, t-test PIX-CON: p = 0.03, Fig 3A) and decreased about 70% in the CBX condition (synchrony = 0.008 ± 0.004, t-test CBX-CON: p = 0.04) compared to the CON condition (synchrony = 0.025 ± 0.018). In addition, when plotting the synchrony as a function of effective coupling averaged for each animal, we found a significant correlation, as expected (regression model in Wilkinson notation (Wilkinson and Rogers, 1973): *synchrony* ∼ 1 + *g*_*eff*_, R^2^ = 0.22, F-test: p = 0.04, Fig 3B) with a positive coefficient (*mean ± sem*, 5 ± 2).

**Figure 3:**
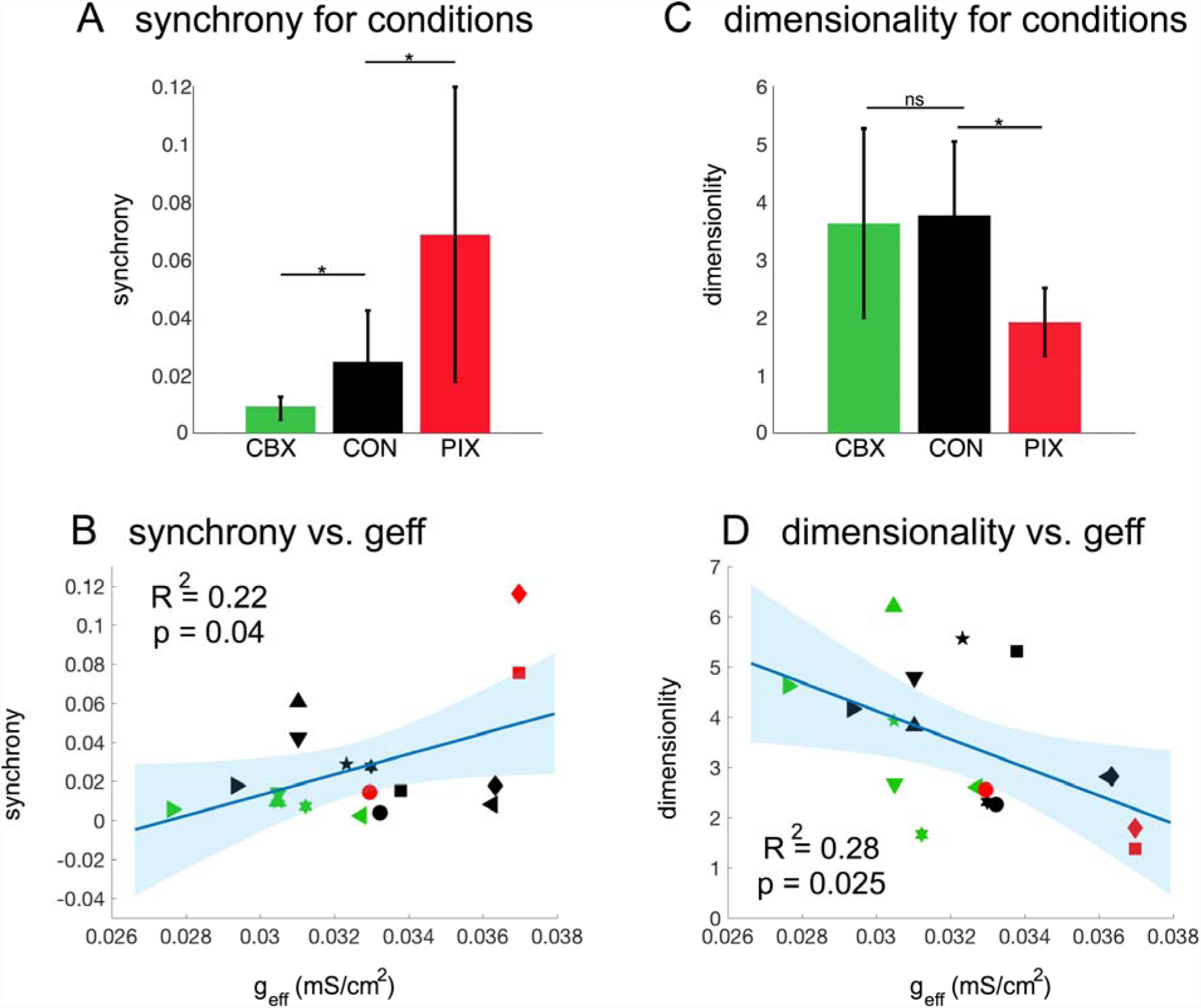
The synchrony and the dimensionality in IO firings moderated by effective coupling. A-C: The synchrony estimated as coincidence of spikes in 1 ms bins (A) and estimation of dimensionality of IO firings (C) for 9 animals in the three data conditions. Error bars are STDs. Asterisks represent significance levels of t-tests: ns p > 0.05, *p < 0.05. B-D: The synchrony (B) and the dimensionality (D) as functions of effective coupling strength averaged for selected neurons in individual animals confirming that effective coupling is a control parameter to optimize the synchrony and thus the dimensionality of IO firings. Each type of symbol represents the data of an individual animal. The cyan solid lines show results of the linear regression models and shaded regions are of 95% CIs.

Next, we estimated the dimensionality (*d*) of complex spike activity for selected neurons in each animal. Briefly, we extracted the average firing rates of neurons in 50-second long periods and applied principal component analysis (PCA) to compute the covariance of these firing rate vectors. *d* has been approximated (by Equation 5) as the minimal number of components accounting for 90% of variability in the data (Abbott et al., 2011). However, with this method, the dimensionality *d* is under-estimated for data with small numbers of neurons, as in our data (Mazzucato et al., 2015). We thus developed a new extrapolation method to correct the dimensionality estimation (see Experimental Procedures for more details). Results showed that *d* was significantly reduced by PIX (*d* = 1.8 ± 0.8, t-test PIX-CON: p = 0.02, Fig 3C) compared to the CON condition (*d* = 3.7 ± 1.2), but there was no significant difference between the CBX (*d* = 3.5 ± 1.6, t-test CBX-CON: p = 0.7) and the CON conditions, probably because of a relatively small effect CBX had on complex spike firing rate. In addition, a regression analysis showed a strong correlation between *g*_*eff*_ and *d* (regression model: *d* ∼ 1 + *g*_*eff*_, R^2^ = 0.28, F-test: p = 0.025, Fig 3D). Here the regression coefficient was negative (*mean ± sem*, −280 ± 113) and opposite to that of synchrony vs. *g*_*eff*_ (*mean ± sem*, 5 ± 2, see above), supporting our hypothesis that synchronization is a feasible mechanism for dimensionality reduction in IO neurons and that effective coupling is the control parameter for the IO to optimize the dimensionality of the olivo-cerebellar system.

### Inverted U-shaped relationship between complexity entropy and effective coupling

Finally, we addressed the question of whether physiological and intermediate coupling strengths maximize the chaotic level of IO activity. The Lyapunov exponents quantify the sensitivity of a dynamical system to initial conditions (Farmer and Sidorowich, 1987; Sano and Sawada, 1985), and are thus often used as indicators of chaos. However, methods to compute Lyapunov exponents from time series data (Kantz, 1994; Rosenstein et al., 1993) are not applicable to our spike data sets, because the computation requires access to continuous variables. We therefore computed the complexity entropy (Letellier 2006; Hirata and Aihara, 2009, see Experimental Procedures for details), which has been shown to approximate the first Lyapunov exponents in simulations of the IO neurons (see Figure S6).

For both the simulated IO spike and the experimental complex spike data sets, we investigated whether the relationship between complexity entropy and effective coupling formed an inverted U-shape, as previously shown in simulations (Schweighofer et al., 2004; Tokuda et al., 2010). For each of the experimental IO neurons, we computed the complexity entropy from the simulated spike data that was generated with the estimated coupling values that best fit the data in terms of the PCA error (difference between experimental and simulated spike data in the PCA space, see Figure S2A). For the IO model, the second order model *entropy ∼ 1 + g*_*eff*_*+ g*_*eff*_^*2*^, where *entropy* is the complexity entropy, had a negative coefficient of the second order term (*mean ± sem*, −246 ± 46), and better fit the simulated spikes in the three conditions than the first-order linear model (*entropy ∼ 1 + g*_*eff*_) (Figure 4A, the Log likelihood ratio (LLR): p < 0.0001). For the IO data (Figure 4B), a mixed effect regression model analysis, with *Animal* as a random intercept accounting for repeated measures within the same animal, showed that the second order model *entropy ∼ 1 + g*_*eff*_*+ g*_*eff*_^2^ *+ (1 | Animal),* where *(1 | Animal)* is the random intercept, had a negative fixed-effect coefficient of the second order term (*mean ± sem*, −217 ± 63), and provided a better fit than the linear model (*entropy ∼ 1 + g*_*eff*_ *+ (1 | Animal),* LLR: p = 0.0007). Thus, for both the IO model (Figure 4A) and the experimental data (Figure 4B), an inverted U-curve that peaks at around *g*_*eff*_ = 0.032 was found, indicating that similar intermediate coupling strengths induce chaotic behavior in both the model and the experimental data. It should be noted that the relatively small changes in the complexity entropy that we observed in the model and data induce significant changes in firing dynamics, from synchronous and rhythmic firings (*entropy* = 0.24, λ_1_ = 20 bits/second, cf. Figure S6B) to chaotic firings (*entropy* = 0.29, λ_1_ = 50 bits/second).

**Figure 4:**
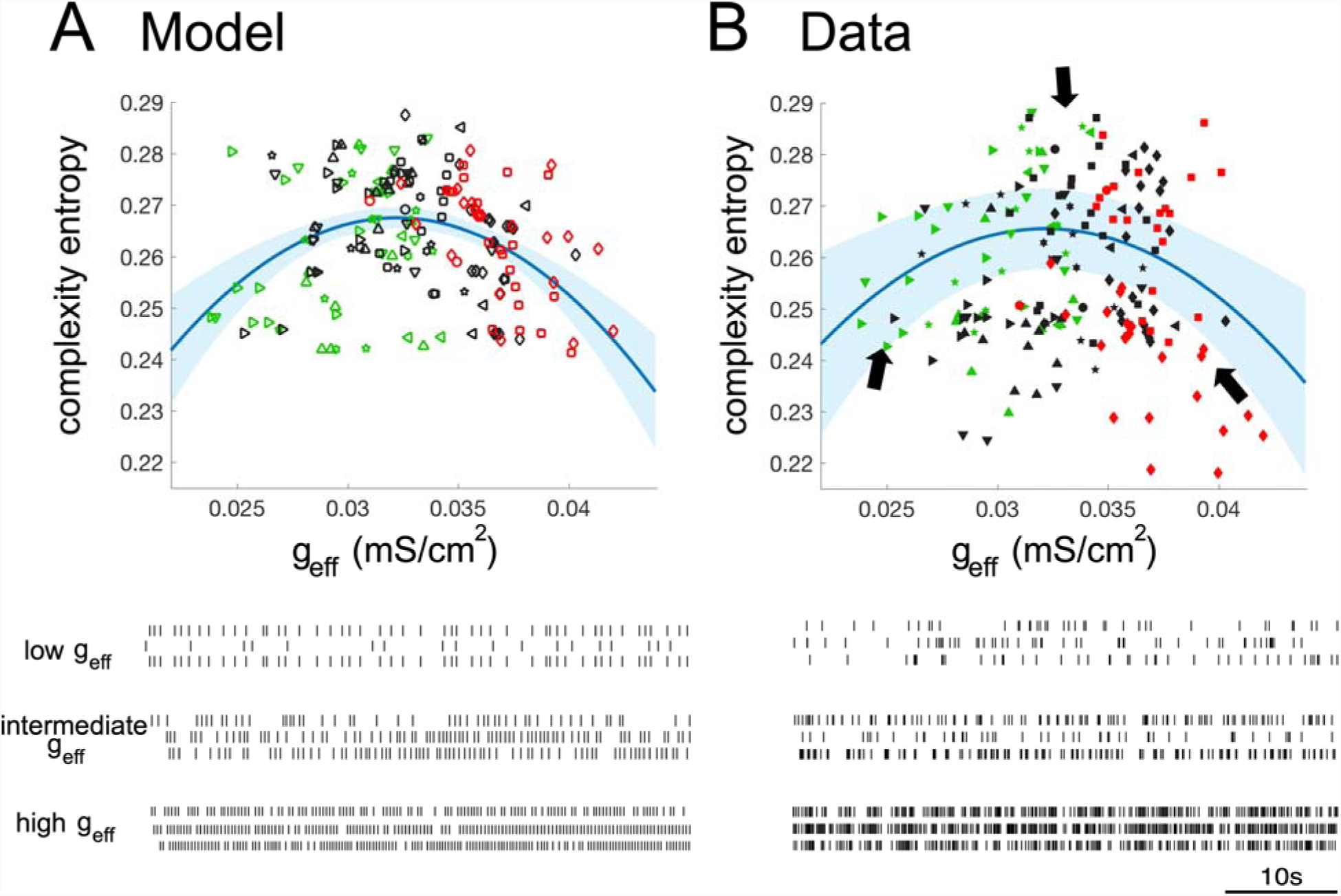
Inverted U-shaped relationship of the complexity versus effective coupling. A-B: Complexity entropy versus effective coupling. Upper panel: chaotic levels measured by the complexity entropy of the spike data as a function of effective coupling strength for the model (A) and real inferior olive neurons (B) confirming that moderate couplings induce chaos. Each value in the model (open symbols) is of the model neuron that best fits to the actual IO neuron in terms of the PCA error. Each type of symbol represents the data of an individual animal. The cyan solid lines indicate the second-order of linear model (A) and mixed-effects model (B) and shaded regions are of 95% CIs. Lower panel: spike trains of the representative neurons (located at dark arrows in the upper right plot) which show periodic and synchronous firings for either low or high couplings but exhibits chaotic firings for intermediate couplings.

## DISCUSSION

We developed a novel technique that combines computational modeling, Bayesian inference and model-averaging to estimate the effective coupling from rat *in vivo* complex spike data. The estimated effective coupling strengths of the three data conditions were consistent with the physiological effects of the drugs, i.e., increased by PIX and decreased by CBX. Notably, the coupling coefficients estimated from our simulations were highly compatible with *in vitro* values observed in (Lefler et al., 2014). In both studies, an approximate doubling of the CC was found for the conditions where GABA activity was lower or blocked. Such compatibility validates the estimation of coupling strengths in our study.

Our analysis of complex-spike data shows that increased electrical coupling between IO neurons decreases the dimensionality of the IO firing dynamics. Dimensionality reduction has long been considered one of the core computations in the brain (Pillow et al., 2008; Cunningham and Yu, 2014; Churchland et al. 2010; Rigotti et al., 2013; Mazzucato et al., 2016). Our study provides direct evidence that electrical coupling among neurons can control the dimensionality of the population activity by modulating the synchrony of the neural code. No significant difference of dimensionality between CBX and CON conditions was found, probably because of the incomplete effect of CBX in de-synchronizing complex spike activity (Blenkinsop and Lang, 2006). Quantitatively, the approximately two-fold reduction in dimensionality from the PIX to the CON condition was highly comparable to that of stimuli-evoked activity of cortical neurons under different stimulus conditions and in varied tasks (Mazzucato et al., 2016; Churchland et al. 2010). We note, however, that additional mechanisms could work in parallel to effectively control the DOFs, such as pruning of irrelevant inputs (Cortese et al. 2018). In the olivo-cerebellar system, in particular, climbing fiber-Purkinje-cell synapses are gradually eliminated based on IO activity during development (Schweighofer, 1998; Good et al., 2017). Further experimental and computational analyses are required to elucidate the interplay between possible mechanisms in controlling the DOFs of the olivo-cerebellar system.

Our results also show that intermediate ranges of electrical coupling induce chaotic dynamics. In contrast, weak or strong coupling decrease the complexity entropy. The finding of an inverted U curve of complexity entropy as a function of effective coupling in both the model and experimental data are consistent with the “chaotic resonance” hypothesis, according to which chaotic firing increases information transmission despite the low firing rates of IO neurons (Schweighofer et al., 2004). We have previously proposed, and shown in simulations, that such chaotic firing may be useful to enhance cerebellar learning by increasing the error transmission capability of the olivocerebellar system (Tokuda et al., 2010). In agreement with this view, a previous study showed that the entropy of neural activity and mutual information between stimulus and response are maximized in balanced excitatory/inhibitory cortical networks (Shew et al., 2011).

Experimental data supporting the importance of electrical coupling for cerebellar learning comes from mice mutants which, due to lacking of electrotonic coupling between IO cells, exhibit deficits in learning-dependent motor tasks (Van Der Giessen et al., 2008). Similarly, humans with reduced IO coupling show motor learning impairments (Van Essen et al., 2010). Because the inhibitory neurons controlling the strength of coupling between IO cells are largely located in the deep cerebellar nuclei (Fredette and Mugnaini, 1991; De Zeeuw et al., 1989), the major output station of the cerebellum, the strength of effective coupling, and thus the level of chaotic behavior, presumably depends on the modulation of the deep cerebellar nuclear neurons via plastic processes in the cerebellar cortex and nuclei (Best and Regehr, 2009; Chaumont et al., 2013; DeGruijl et al., 2014; Lefler et al., 2014; Turecek et al., 2014; Witter et al., 2013). Indeed, changes in simple spike levels produce significant changes in complex spike synchrony (Marshall and Lang, 2009). Thus, the IO coupling strength during cerebellar learning could be adaptively modulated, with the Purkinje cell-cerebellar nuclei-inferior olive triangle acting to decrease coupling along the progress of cerebellar learning (Kawato et al., 2011; Schweighofer et al., 2013; Tokuda et al., 2013).

According to this view, in the early phase of learning, the motor commands are strongly disturbed and far from the desired ones. The Purkinje cells, which are then strongly modulated by large sensory inputs and error signals, suppress the inhibitory effect of the neurons in cerebellar nuclei on the IO. Thus, the IO neurons are initially strongly coupled and the dimensionality is low. Because of this low dimensionality, the IO network would respond only to low-dimensional components of the error signals, which may convey only the gross features of the motor commands. However, the strong coupling allows a widespread synchrony among IO neurons and potentially leads to massive changes in the parallel-fiber-Purkinje-cell synaptic weights, resulting in fast but coarse learning. In addition to its effect on learning, such highly synchronized IO activity may have a downstream effect via a large network of synchronized Purkinje cells (Blenkinsop and Lang, 2011; Lang and Blenkinsop 2011; Tang et al., 2016), that could trigger an emergency or protective feedback motor commands in response to this error. In contrast, in the late phase of learning, as the motor error becomes smaller, the Purkinje cell activity may become weaker, allowing increased activity of cerebellar nuclear neurons. This would result, in turn, in reduced IO coupling and higher dimensionality. At this stage, the moderate coupling strengths could induce chaotic IO spike activity that would transmit high dimensional error signals, resulting in more sophisticated learning (Shaikh et al., 2017). A further possibility is that these high dimensional signals would also be used for the fine grain motor control commands that are needed for precise motor coordination (Hoogland et al, 2015).

## ACKNOWLEDGEMENTS

HH, KT and MK were supported by a contract with the National Institute of Information and Communications Technology entitled ‘Development of network dynamics modeling methods for human brain data simulation systems’. HH and KT are partially supported by ImPACT Program of Council of Science, Technology and Innovation (Cabinet Office, Government of Japan). HH, ITT and KT are partially supported by MEXT Kakenhi (No. 17H06313). HH is partially supported by JST ERATO (JPMJER1801, “Brain-AI hybrid”). KA is partially supported by MEXT Kakenhi (No. 15H05707) and AMED under Grant Number JP18dm0307009. EJL acknowledges grants NIH NS095089 and NS37028. NS acknowledges grants NSF BCS-1031899 and 1R56NS100528-01.

## EXPERIMENTAL PROCEDURES

The recording experiments were performed in accordance with the National Institute of Health Guide for the Care and Use of Laboratory Animals. Experimental protocols were approved by the Institutional Animal Care and Use Committee of New York University School of Medicine.

### Experimental data

The analyses were performed on a subset of data obtained in two prior series of experiments in ketamine/xylazine anesthetized female, Sprague-Dawley rats that involved either injection of picrotoxin (PIX) or carbenoxolone (CBX) to the IO to block GABA-A receptors or gap junctions, respectively (Blenkinsop and Lang, 2006; Lang, 2002; Lang et al., 1996). The specific experiments were chosen primarily on the basis of having typical complex spike activity in control and a large change in activity in response to the drug injection.

Details of the experimental procedures can be found the original reports. In brief, a rectangular array of glass microelectrodes was implanted into the apical surface of crus 2a. The arrays typically contained 3-4 mediolaterally running rows and up to 10 rostrocaudally running columns, with an interelectrode spacing of ∼250 μm. Electrodes were implanted to a depth of ∼100 μm below the brain surface such that complex spikes from individual Purkinje cells were recorded. In each experiment, spontaneous complex spike activity was recorded during an initial control period. Following the control (CON) period, the IO was located by lowering a microelectrode through the brainstem under stereotaxic guidance until activity characteristic of IO neurons was observed. The microelectrode was then replaced by an injection pipette containing the drug solution that was lowered to the same location as the site where IO activity was found. A slow injection of drug solution was then performed (∼1 μl over 5-10 min). The drug conditions analyzed were recorded after completion of the injection and a clear change in activity was observed. The multielectrode arrays recorded from 10 - 30 Purkinje cells in each of the CBX experiments (n = 6 animals), and from 16-42 Purkinje cells in the PIX experiments (n = 3 animals).

The effect of CBX and PIX on complex spike activity often varied among cells within an experiment. This was likely due to the Purkinje cells in different parts of the array receiving climbing fibers from different regions of the IO, that the drugs were injected at a single point within the IO, and that drug concentration (and therefore potentially efficacy) will fall with distance from the injection site. Indeed, the IO is an extended structure (particularly in the rostrocaudal axis where it is ∼2 mm long). We therefore considered the effects of the drugs when selecting the neurons for analysis. That is, Purkinje cells that exhibited significant changes in complex spike synchrony, measured as the coincidence of spikes in 1 ms time bins, between the control and drug conditions were selected. For CBX, the criterion was a 50% decrease and for PIX it was a 200% increase. In total, we analyzed spike train data from 500-second long periods for the control and drug conditions for each neuron (neurons/condition: control, n = 90; PIX, n = 46; CBX, n= 44).

### IO network model

The IO neuron model is a conductance-based model (Schweighofer et al., 1999) extended via addition of glomerular compartments comprising electrically coupled spines (Onizuka et al., 2013). The network model consisted of an array of 3×3 IO neurons, each of which was mutually connected to its four neighboring neurons by a gap junction from one of its spines to one of its neighbor’s represented by the gap-junctional conductance *g*_*c*_. We simulated spike data of the nine cells with step-wise changes of two model parameters, i.e., inhibitory synaptic conductance *g*_*i*_, and coupling conductance *g*_*c*_. These two parameters were both varied in the range of 0–2.0 mS/cm^2^ with an increment of 0.05 mS/cm^2^. We generated a total of 41×41=1681 sets of 500-second long simulated spike trains. The simulated spike data for each variation of *g*_*i*_ and *g*_*c*_ was then compared with the actual spike data, and the parameters whose firing dynamics best fit to that of individual neurons in the control, PIX, and CBX conditions were selected as the estimated values. Because the effect of the axial conductance of the spines, *g*_*s*_, is equivalent to that of the gap-junctional conductance, *g*_*c*_, in determining the amount of current will flow across the gap junction, *g*_*s*_ does not need to be estimated from the data and thus was fixed at 0.1 mS/cm^2^ (Onizuka et al., 2013). To better account for excitability of the neurons *in vivo*, the inward sodium current conductance *g*_*Na*_ was set as 110 mS/cm^2^, which has been shown to induce robust chaos in the model (Schweighofer et al., 2004).

### The segmental Bayes inference for estimating the effective coupling from a single model

Under simplified assumptions, the effective coupling, *g*_*eff*_, between two IO neurons was calculated from the axial conductance of the spines *g*_*s*_, inhibitory conductance *g*_*i*_ and gap-junctional conductance *g*_*c*_ as in (Katori et al., 2010):

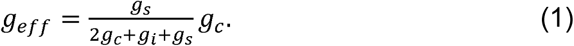

This equation implies that to estimate the effective coupling *g*_*eff*_, we need to estimate both the coupling conductance *g*_*c*_ and the GABA conductance *g*_*i*_ reliably for each of the three datasets CBX, CON, and PIX. For that purpose, we previously developed a Bayesian method that contains two steps (see Supplemental Materials for outlines of the Bayesian method, and Hoang et al., 2015). In the first step, the parameters are estimated for each 50-second time-segment of individual neurons, allowing the parameter values to vary in time. This compensates for inevitable mismatch in the firing patterns between the model and the data. In the second step, a single set of parameter values is estimated for the entire time-segments of individual neurons by a hierarchical Bayes framework. Below, we outline the segmental Bayes method (for a detailed description see, Hoang et al., 2015).

First, the firing dynamics of the spike data were characterized by a feature vector composed of a total of sixty-eight spatiotemporal features, e.g., firing rate, local variation (Shinomoto et al., 2005), cross-correlation, auto-correlation, and minimal distance (Hirata and Aihara, 2009). Principal component analysis (PCA) was then conducted to remove the redundancy of those features. The Bayesian inference aims to inversely estimate the conductance values from the top three-dimensional principal components, which accounted for 55% of the data variance. To compensate for the modeling errors, i.e. differences in the complexity of firing patterns between the model and actual neurons, we divided the spike data of each neuron into short time-segments under the assumption that segmental estimates of individual neurons fluctuated around a single neuronal estimate with a normal (Gaussian) distribution. The conductance values of individual neurons can be estimated by a hierarchical Bayesian framework. Here, the segment size, 50 seconds, was optimized so that the variance of firing frequency across segments was minimal (Onizuka et al., 2013). In addition, we also introduced two physiological constraints on the estimates: a common *g*_*i*_ for CBX and CON neurons in CBX experiments and a common *g*_*c*_ for PIX and CON neurons in PIX experiments. The rationale for these constraints is that CBX and PIX are supposed to only reduce the gap-junctional conductance *g*_*c*_ and inhibitory conductance *g*_*i*_, respectively.

We have shown that the segmental Bayes algorithm minimizes the fitting between experimental and simulated spike data (Hoang et al., 2015), and further confirmed, by simulations, that it indeed minimizes the estimation errors compared to other conventional methods – including the non-segmental Bayes inference, which finds the estimates once across the entire spike data, and the minimum-error algorithm, which directly finds the closest match in the feature space (data not shown).

### Model-averaging estimation of the effective coupling

We found that the firing frequency of inhibitory synaptic noise inputs largely affect spiking behavior of the IO model and thus the estimation results. To reduce the uncertainty in estimates of *g*_*c*_ and *g*_*i*_, we therefore adopted the segmental Bayes algorithm by a model-averaging approach as follows (for review, see Grueber et al., 2011). Due to the extremely expensive computation of the compartmental model (about a week for ten computer clusters to generate the spiking data of 1681 conductance values of a single model), we first simulated four models with the firing frequency of inhibitory synaptic inputs of 10, 20, 50 and 70 Hz, which are observed in slices of cerebellar nucleo-olivary neurons (Najac and Raman, 2015). Next, we conducted the segmental Bayes to estimate posterior probability of *g*_*i*_ and *g*_*c*_ for each model.

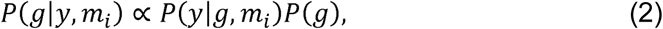

where *P(g | y, m*_*i*_) is the posterior probability of the conductance *g = (g*_*i*_*, g*_*c*_), *y* is the feature vectors extracted from the spike data, and *m*_*i*_ is the *i*th selected model (*i* = 1‥4). We then mixed the posterior probabilities with the weights proportional to the model evidence as follows:

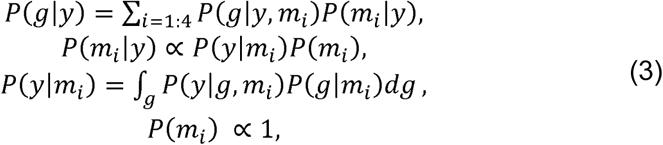

where *P(g | y)* is the mixed probability for an individual neuron and *P(y | m*_*i*_) is the evidence of the *i*th model. Here, all models are treated equally with a non-preference prior *P(m*_*i*_). Finally, the point estimates of *g*_*i*_ and *g*_*c*_ were computed by marginalizing the mixed posterior probabilities.

### Calculation of the synchrony for individual neurons

The spike train of a neuron was binned into *X(i)*, where *i* represents the time step *(i = 1,…,T)*, with *X(i) = 1* if the spike occurs in the *i*th time bin; otherwise, *X(i) = 0*. The synchrony of two different neurons, *x* and *y*, was calculated as the cross-correlation coefficient at zero-time lag:

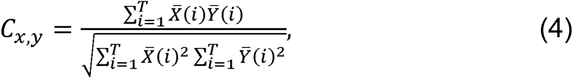

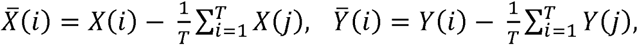

where 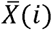 and 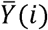 are the normalized forms of *X(i)* and *Y(i)* to account for the firing frequency. Here, the two spikes were considered synchronous if their onsets occur in the same 1 ms bin. The synchrony level of an individual neuron *x* was computed as the mean of *C*_*x,y*_ for all neurons *y*≠*x* in the same animal.

### Estimation of the dimensionality of neural firings

The dimensionality can be considered as the minimal dimensions necessary to provide accurate description of neural dynamics. The principal component analysis (PCA) has become the most widely used approach because it enables to represent neural dynamics in a lower dimensional space (Mazzucato et al. 2016). Here, we adopted this approach for estimating the dimensionality of the IO firing activity in the presence of a small number of recorded neurons.

We first segmented the IO spike trains into time windows of 50 seconds, from which the firing rate vectors of all neurons were computed (see Fig S5A). Firing rate vector in each sampled window corresponds to an observation in the *N*-dimensional space, where *N* is the number of ensemble neurons. Then, PCA was applied to estimate the dimensionality as (Abbott et al. 2011):

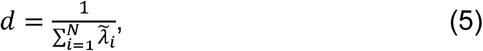

where 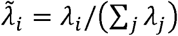 are the principal eigenvalues expressed as the amount of variance explained (see Fig S5A), and *λ*_*i*_ is the *i*th eigenvalue of the covariance matrix of the firing rate vectors.

It is noted that the dimensionality was not sensitive to the length of sampled window (10-50 seconds were analyzed but no significant different values were found, Fig S5B) probably because the IO firings are rather stable across the time course. However, it has been shown that dimensionality estimation depends on the number of ensemble neurons *N*. Specifically, d is underestimated for small *N* but becomes independent of *N* for sufficiently large *N* (Mazzucato et al., 2016). After data selection (see above), the number of IO neurons in each animal is n = 3-22, which is likely to suffer from the under-sampling bias. To overcome such challenge, we computed the corrected values of dimensionality following the quadratic extrapolation method (Shew et al., 2011). First, we randomly selected a fraction *f* of *N* neurons. We then computed the dimensionality *d*, follows the Equation 5, for fractions *f* = 0.2 to 1 in steps of 0.2, repeated 50 times for each *f*. Next, the average *d* versus *f* data was fitted by the following model (Fig S5C):

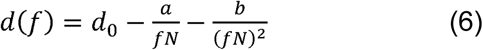

The fit parameter *d*_*0*_ is the corrected estimate of the dimensionality and is reported in the text.

### Computation of the complexity entropy

The Lyapunov exponents quantify the sensitivity of a dynamical system to initial conditions, and thus are often used as indicators of chaos (Farmer and Sidorowich, 1987; Sano and Sawada, 1985). A number of methods have been developed to compute the Lyapunov exponents from time series with a fixed sampling interval (Kantz, 1994; Rosenstein et al., 1993). Those methods, however, are not applicable for our IO data because computation of Lyapunov exponents requires access to continuous variables, which is not the case in our discrete IO spike sets. We therefore adopted a previously proposed approach (Hirata et al., 2008) that approximates the Lyapunov exponents via a recurrence plot by using the edit distance of spike trains (Victor and Purpura, 1997). Our method requires computing the modified edit distance of the spike trains (Hirata and Aihara, 2009) and its recurrence plot (Eckmann et al., 1987; Marwan et al., 2007). The complexity entropy (Letellier, 2006) was computed from the distribution of the length of diagonal lines in the recurrence plot (see Figure S6 for illustration of the complexity method).

We first sampled the spikes trains in windows of 20 seconds and computed the edit distance for all pairs of sampled windows. To resolve the issue of discontinuity induced by the difference in the number of spikes in two sampled windows, we adopted a modified version of edit distance computation as in Hirata and Aihara (2009). Briefly, for each sampled window, we took into account the spikes that occur immediately before and/or after the time window, thus resulting in four derived windows. We then computed the edit distance for a total of 16 (4×4) derived pairs of the two sampled windows and temporarily assigned the minimum value as edit distance between them. The edit distance of two derived windows is defined by a total minimal cost for converting one window to the other (Victor and Purpura, 1997). Allowed operations include deletion or insertion of events (both cost 1 for each event), and shift of events (cost 20% the amount of shifting in second for each event). The edit distance for all pairs of sampled windows of 20 seconds with an interval of 2 seconds constitutes a two-dimensional distance matrix. We then updated the edit distance matrix by the shortest distance connecting any two sampled windows – see Figure S6A.

The recurrence plot is constructed by binarizing the edit distance matrix, with the distance values smaller than a predefined threshold as 1, and the others else as 0 (Eckmann et al., 1987). The threshold was determined so that 30% of data points in the distance matrix were 1, as in (Marwan et al., 2007). Next, we extracted the frequency distribution of the length of the points 1 that form diagonal lines in the recurrence plot. The Shannon entropy of that distribution has been shown to be inversely proportional to the largest Lyapunov exponent (Letellier, 2006). We thus used the inverse of Shannon entropy as a measure of chaos for the spike data.

To validate that complexity entropy is an indicator of chaos, we generated noise-free simulation data and computed the correlations between complexity entropy and the Lyapunov indexes (c.f. Figure S6B-C). Note that this approach is possible for the simulation data because we have access to the continuous trace of the membrane potential. Specifically, we first removed the noise in the synaptic inputs, and simulated 500-second spike trains for more than 100 conductance values (*g*_*i*_ varied in 0–1.0 mS/cm^2^ and *g*_*c*_ in 0–2.0 mS/cm^2^) and estimated the complexity entropy from the simulated spike trains. Next we computed the Lyapunov exponents of the IO model by the method of (Wolf et al., 1985), and then extracted the largest component, λ_1_, as well as the Lyapunov dimension, *D*_*KY*_, as these are two direct indicators of chaos (Kaplan and Yorke, 1970).

## Statistical Analysis

Unless specifically stated elsewhere, all data is reported as *mean* ± *std*. The non-parametric Kruskal-Wallis one-way analysis of variance was used to test whether data groups of different sizes originate from the same distribution.

## SUPPLEMENTAL INFORMATION

**Figure S1:**
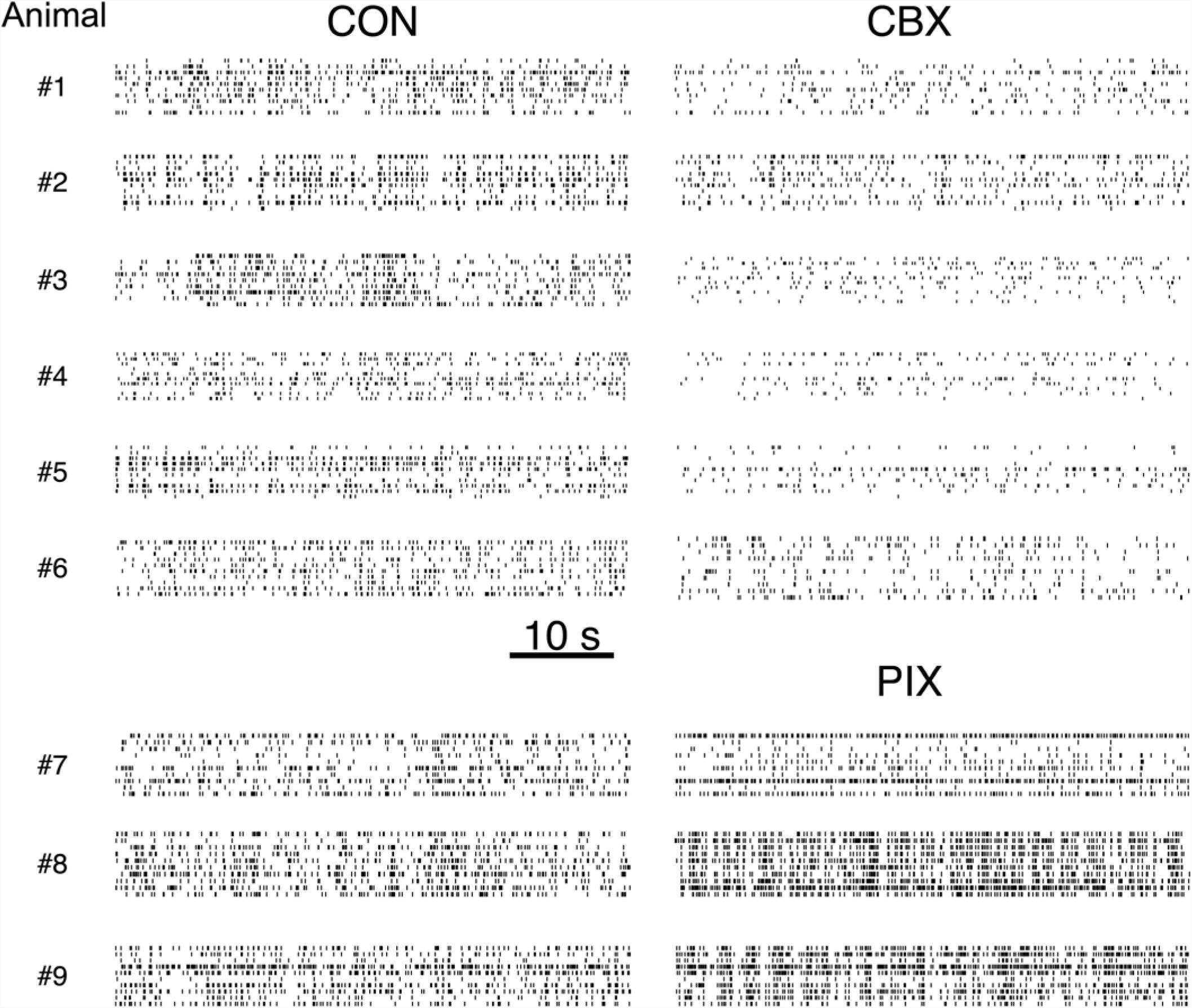
Inferior olive firing data set for all animals. A: Spike data in 50 second of 10 representative neurons in 9 animals with the physiological conditions (CBX and PIX) in the right and the control condition (CON) in the left columns.

**Figure S2:**
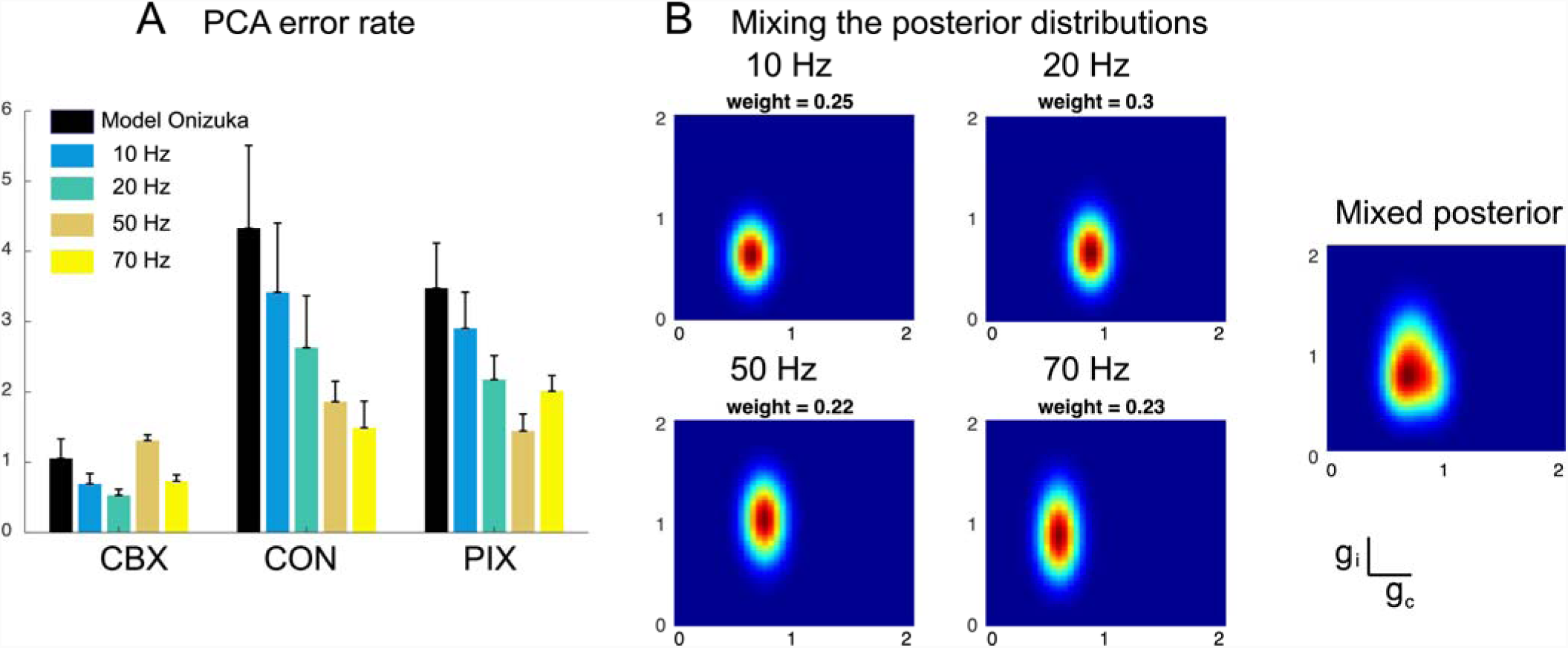
Improving the parameter estimates via Bayesian model-averaging. A: PCA error rates of the *g*_*i*_ and *g*_*c*_ estimates by the segmental Bayesian inference averaged for the entire IO neurons for CBX, CON, and PIX conditions for four different models (color bars) in comparison with the previous model (black bar, Onizuka et al., 2013). The error bars are of 95% CIs. B: Posterior probabilities of a representative IO neuron by individual models and the mixed posterior probability with the weights determined by the evidence of Bayesian inference.

**Figure S3:**
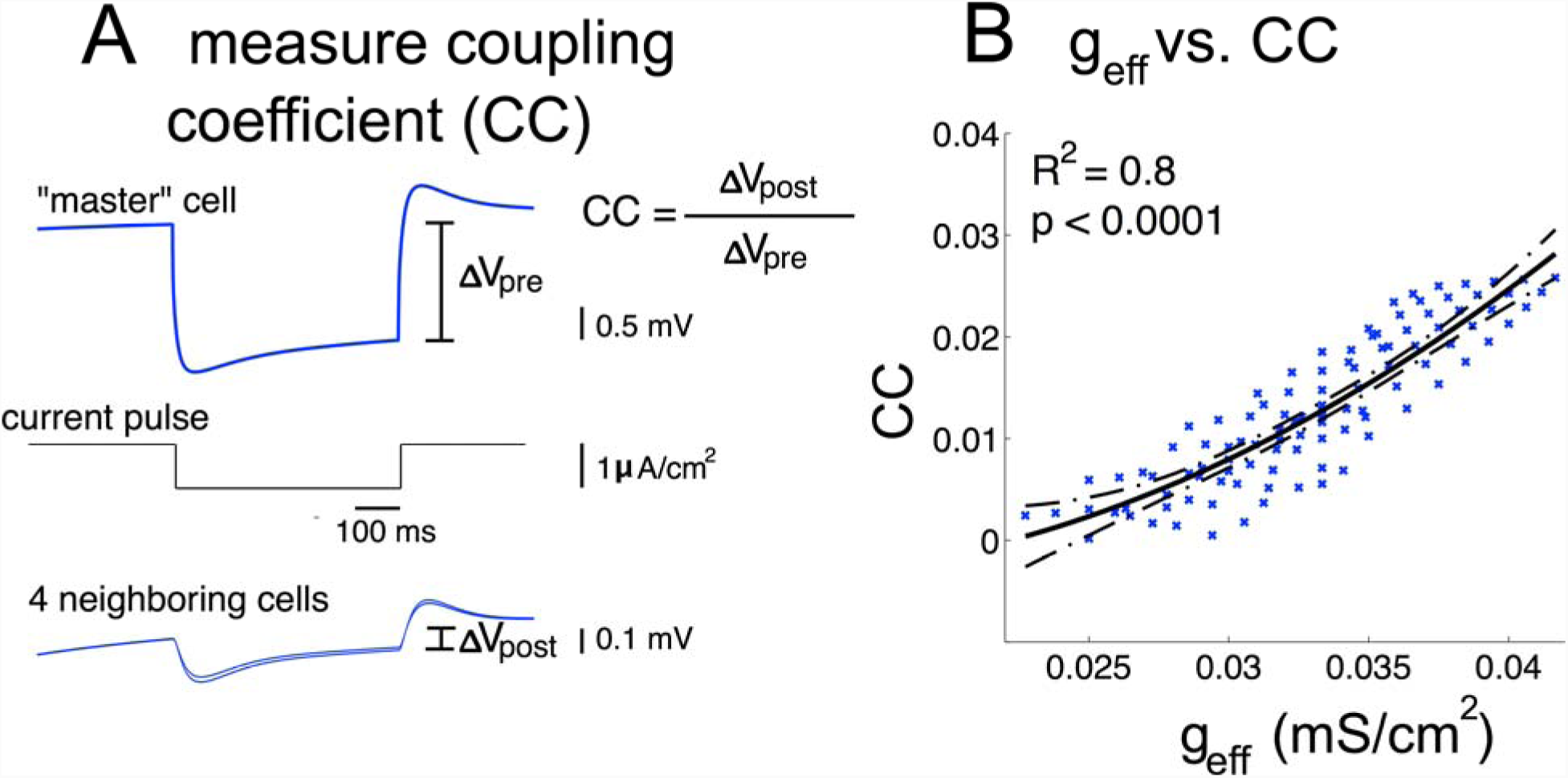
Estimation of the coupling coefficient (CC) by simulation. We injected a current pulse of −1 μA/cm2 to the cell #5 and recorded the steady-state voltage change of this “master” cell and its four post-junctional cells (left panel). We computed the CCs for hundreds of of *g*_*i*_ and *g*_*c*_ values in the range, in which the estimated conductance of the data distributed, and found a strong positive correlation between the effective coupling and the CC (right panel, R^2^ = 0.8, p < 0.0001). It is noted that the non-linear fit represents the nature of deriving *g*_*eff*_ from *g*_*i*_ and *g*_*c*_ follow the Equation (1).

**Figure S4:**
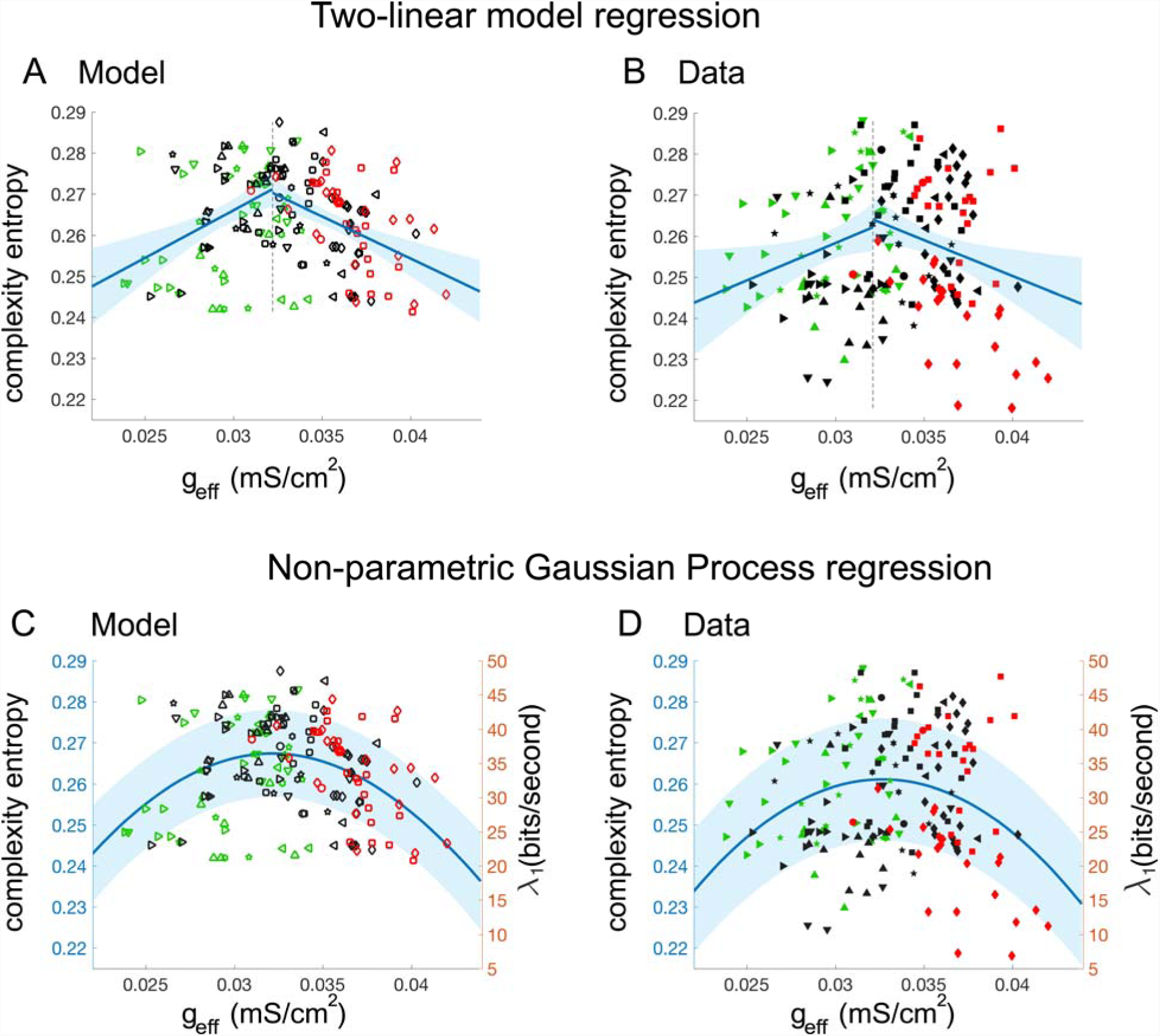
Validation of the inverted U-shaped curves. We investigated whether intermediate couplings maximize the complexity entropy by applying the two-linear regression models as follows. We first divided the data into two separate partitions by the intermediate couplings around the mean *g*_*eff*_ = 0.032 (black dashed lines). Linear regressions were then conducted for the two partitions independently for both the model (A) and the data (B). Results show significant positive coefficients (*mean* ± *sem*, model: 2.3 ± 0.6, p = 0.0002; data: 1.8 ± 0.8, p = 0.03) for the left partitions and significant negative coefficients (*mean* ± *sem*, model: −2 ± 0.4, p < 0.0001; data: −1.7 ± 0.6, p = 0.009) for the right ones. We further applied a non-parametric Gaussian Process regression model, which does not assume an explicit relationship between the coupling and the complexity entropy. Still, we observed inverted U-shaped curves maximized at around *g*_*eff*_ = 0.032 for both the model (C) and the data (D). In sum, these results support the inverted U-shaped relationship between the effective coupling and complexity entropy. The right ordinates of C-D represent the first Lyapunov exponents approximated from the simulation data (c.f Figure S6B), indicating that intermediate couplings induce chaos. The shaded regions in A-B are of 95% CIs and those in C-D are of ±sem.

**Figure S5.**
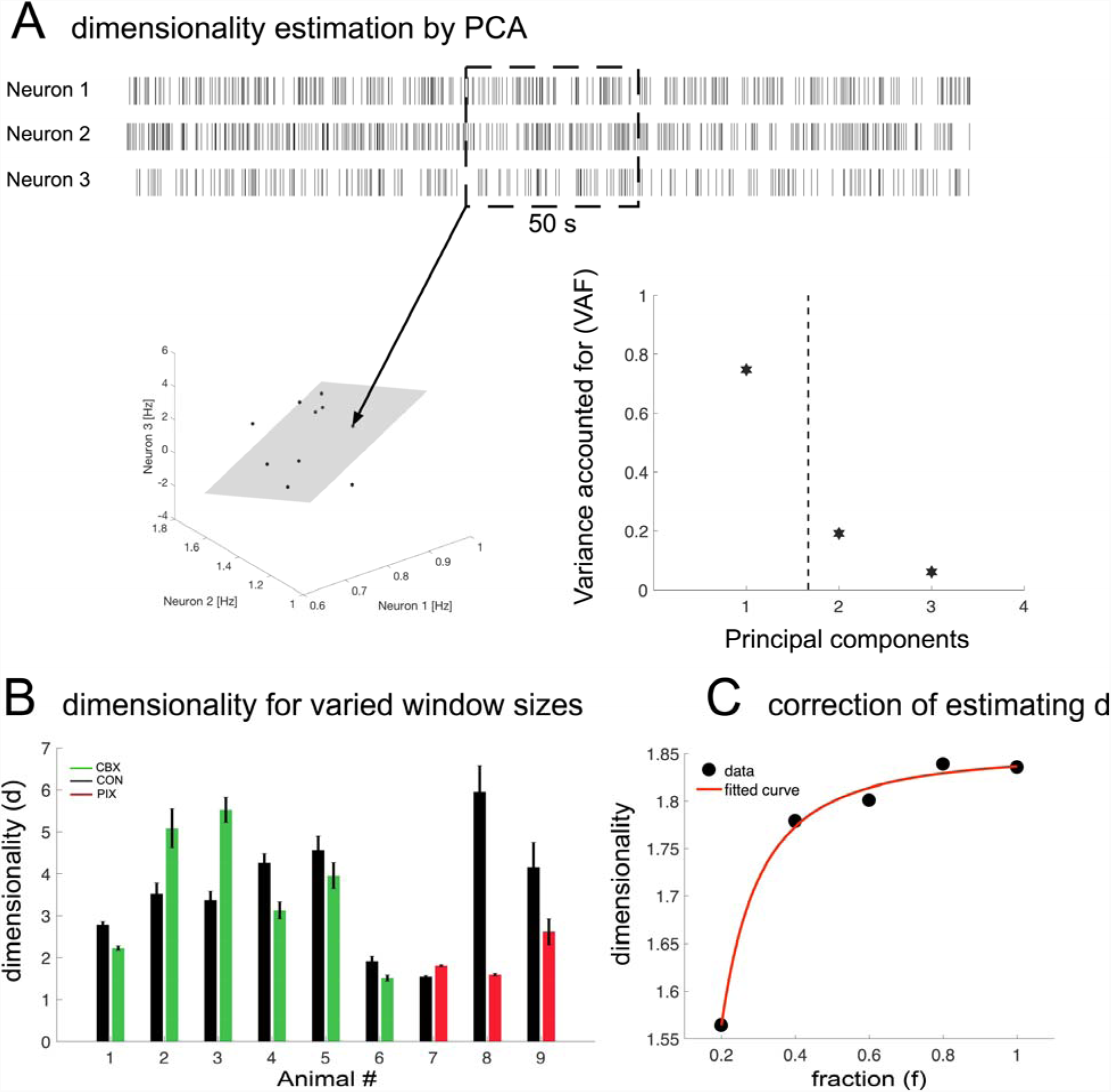
Dimensionality estimation for the spike data of ensemble neurons. A: Illustration of the principal component analysis (PCA) for the firing rate vectors extracted from 50-second windows of three neurons of Animal #6 in the CON condition. The estimated dimensionality *d* = 1.86 (dashed dark line, c.f Equation 5), indicates that the approximately 2-dimensional subspace (shaded gray plane) can explain more than 90% of the variance of neural firing dynamics. B: Estimating dimensionality (Equation 5) with varied window lengths from 10-50 seconds for 9 animals in the three data conditions showing the robustness of dimensionality estimation against the window length. C: under sampling correction of dimensionality estimation for the cases for which the number of recorded neurons is small by quadratic extrapolation fitting of the dimensionality *d* vs. fraction of selected neurons *f*.

**Figure S6.**
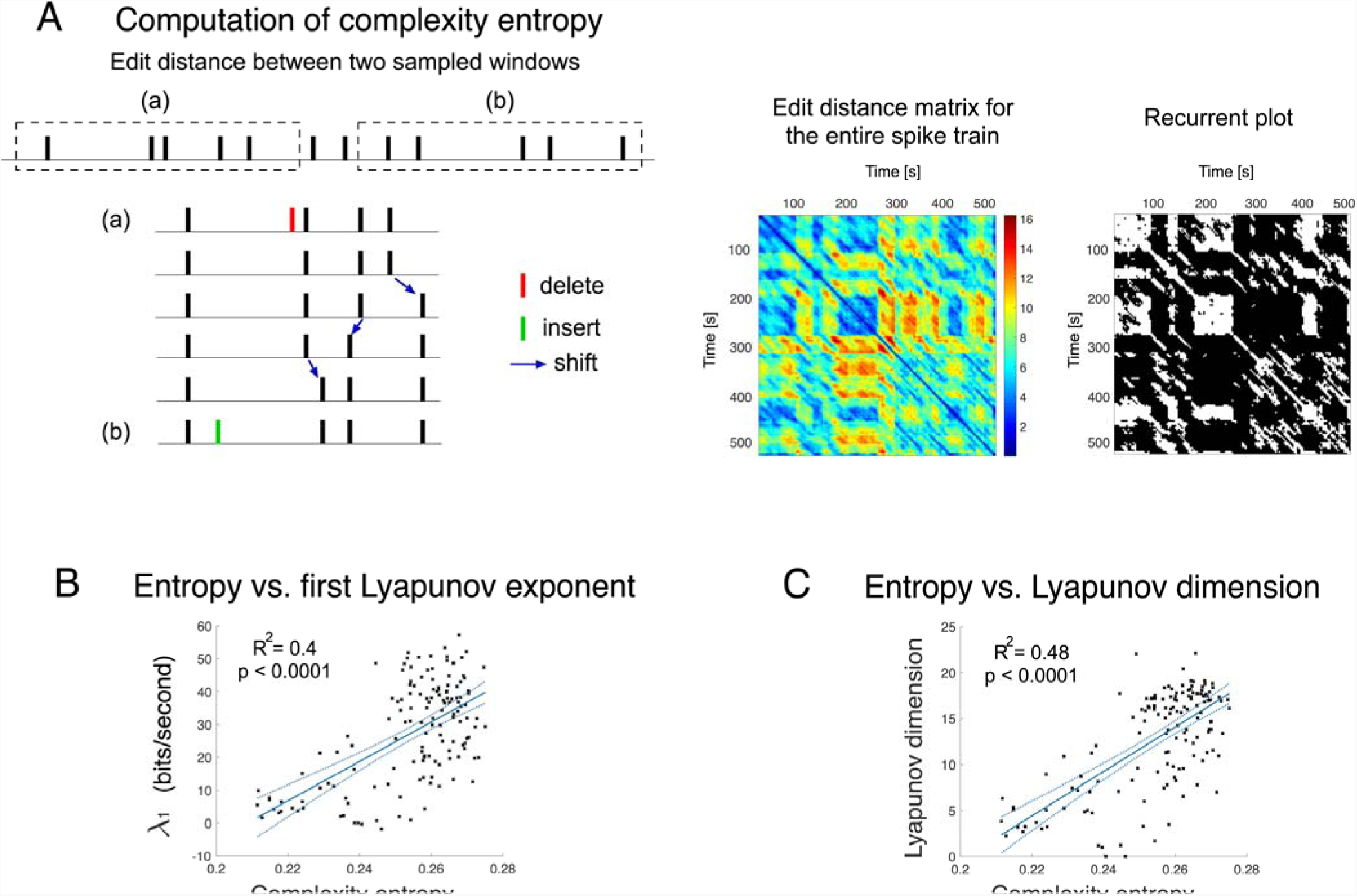
Computation and validation of the complexity entropy method. A: Illustration of edit distance computation between two sampled spike windows shows a sequence of elementary steps that convert the spike window (a) into (b). Each bar represents one spike. Allowed operations include deletion of a spike (shown in red), insertion of a spike (shown in green), or shifting a spike in time (blue arrows). Computation of edit distance for continuous sampling windows for the entire spike train constitutes the edit distance matrix. Then, the recurrent plot is constructed by binarizing the edit distance matrix. The points whose values are smaller than the threshold were plotted as white dots, otherwise as black dots. Complexity entropy is computed as the inverse of Shannon entropy, in terms of frequency distribution of the length of the diagonal lines of white dots (Letellier, 2006). B-C: Complexity entropy measured for a total of a hundred of parameter values (black crosses) in noise-free simulations showed strong positive correlations with the largest Lyapunov exponent λ_1_ (regression model: λ_1_ ∼ 1 + *entropy*, R^2^ = 0.4, F-test: p < 0.0001, Figure S6B) and the Lyapunov dimension D_KY_ (*D*_*KY*_ ∼ 1 + *entropy*, R^2^ = 0.48, F-test: p < 0.0001, Figure S6C). Solid cyan lines represent the fit of linear models with 95% CIs (dashed cyan lines).

## Footnotes

1 Purkinje cell complex spikes, as opposed to simple spikes that are due to the granule cell inputs, bear a one-to-one relationship to IO spikes. Thus, complex spikes can be used as a proxy for IO spikes (see Figure S1 for examples of complex spike recordings in these three conditions; see Methods for experimental procedures).

## REFERENCES

Abbott, L. F., Rajan, K., and Sompolinsky, H. (2011). “Interactions between intrinsic and stimulus-evoked activity in recurrent neural networks,” in The Dynamic Brain: An Exploration of Neuronal Variability and its Functional Significance, eds D. L. Glanzman and M. Ding (New York, NY: Oxford University Press), 65–82.

Adolph KE, Cole WG, Komati M, Garciaguirre JS, Badaly D, Lingeman JM, Chan GLY, Sotsky RB (2012). How do you learn to walk? Thousands of steps and dozens of falls per day. Psychol. Sci., 23:1387–94.

Apps, R. and Hawkes, R. (2009). Cerebellar cortical organization: a one-map hypothesis. Nat Rev Neurosci. 10(9):670–81.

Atkeson CG, Benzun PWB, Banerjee N, Berenson D, Bove CP, Cui X, DeDonato M, Du R, Feng S, Franklin P, et al. (2018). What Happened at the DARPA Robotics Challenge Finals. In The DARPA Robotics Challenge Finals: Humanoid Robots To The Rescue. edited by Spenko M, Buerger S, Iagnemma K. Springer International Publishing: 667–684.

Bastian, A.J. (2006). Learning to predict the future: the cerebellum adapts feedforward movement control. Curr. Opin. Neurobiol. 16, 645–649.

Bazzigaluppi, P., Ruigrok, T., Saisan, P., de Zeeuw, C.I., and de Jeu, M. (2012). Properties of the Nucleo-Olivary Pathway: An In Vivo Whole-Cell Patch Clamp Study. PLoS One 7.

Belluardo, N., Mudò, G., Trovato-Salinaro, A., Le Gurun, S., Charollais, A., Serre-Beinier, V., Amato, G., Haefliger, J.A., Meda, P. and Condorelli D.F. (2000). Expression of connexin36 in the adult and developing rat brain. Brain Res. 865(1):121–38.

Best, A.R., and Regehr, W.G. (2009). Inhibitory Regulation of Electrically Coupled Neurons in the Inferior Olive Is Mediated by Asynchronous Release of GABA. Neuron 62, 555–565.

Blenkinsop, T.A., and Lang, E.J. (2006). Block of Inferior Olive Gap Junctional Coupling Decreases Purkinje Cell Complex Spike Synchrony and Rhythmicity. J. Neurosci. 26, 1739–1748.

Blenkinsop, T. A., and Lang, E. J. (2011). Synaptic action of the olivocerebellar system on cerebellar nuclear spike activity. J Neurosci. 31(41), 14708–14720.

Chaumont, J., Guyon, N., Valera, A.M., Dugue, G.P., Popa, D., Marcaggi, P., Gautheron, V., Reibel-Foisset, S., Dieudonne, S., Stephan, A., et al. (2013). Clusters of cerebellar Purkinje cells control their afferent climbing fiber discharge. Proc. Natl. Acad. Sci. 110, 16223–16228.

Churchland MM, Yu BM, Cunningham JP, Sugrue LP, Cohen MR, Corrado GS, Newsome WT, Clark AM, Hosseini P, Scott BB, Bradley DC, Smith MA, Kohn A, Movshon JA, Armstrong KM, Moore T, Chang SW, Snyder LH, Lisberger SG, Priebe NJ, Finn IM, Ferster D, Ryu SI, Santhanam G, Sahani M, Shenoy KV. (2010). Stimulus onset quenches neural variability: a widespread cortical phenomenon. Nat Neurosci. 13(3):369–78

Condorelli, D.F., Parenti, R., Spinella, F., Trovato Salinaro, A., Belluardo, N., Cardile, V. and Cicirata F. (1998). Cloning of a new gap junction gene (Cx36) highly expressed in mammalian brain neurons. Eur J Neurosci. 10(3):1202–8.

Cortese A., De Martino B. and Kawato M. (2018). The neural and cognitive architecture for learning from a small sample, arXiv.org,1810.02476.

Cunningham J.P. & Yu B.M. (2014). Dimensionality reduction for large-scale neural recordings. Nature Neurosci. 17:1500–1509

DAngelo, E., Mapelli, L., Casellato, C., Garrido, J.A., Luque, N., Monaco, J., Prestori, F., Pedrocchi, A., and Ros, E. (2016). Distributed Circuit Plasticity: New Clues for the Cerebellar Mechanisms of Learning. Cerebellum 15, 139–151.

DeGruijl, J.R., Sokól, P.A., Negrello, M., and DeZeeuw, C.I. (2014). Modulation of electrotonic coupling in the inferior olive by inhibitory and excitatory inputs: Integration in the glomerulus. Neuron 81, 1215–1217.

Devor, A., and Yarom, Y. (2002). Electrotonic coupling in the inferior olivary nucleus revealed by simultaneous double patch recordings. J. Neurophysiol. 87, 3048–3058.

Eccles, J.C., Llinás, R., and Sasaki, K. (1966). The excitatory synaptic action of climbing fibres on the purkinje cells of the cerebellum. J. Physiol. 182, 268–296.

Eckmann, J.-P., Kamphorst, S.O., and Ruelle, D. (1987). Recurrence Plots of Dynamical Systems. Europhys. Lett. 4, 973–977.

Van Essen, T.A., Van der Giessen, R.S., Koekkoek, S.K.E., VanderWerf, F., De Zeeuw, C.I., van Genderen, P.J.J., Overbosch, D., and de Jeu, M.T.G. (2010). Anti-malaria drug mefloquine induces motor learning deficits in humans. Front. Neurosci. 4.

Farmer, J.D., and Sidorowich, J.J. (1987). Predicting chaotic time series. Phys. Rev. Lett. 59, 845–848.

Fredette, B.J., and Mugnaini, E. (1991). The GABAergic cerebello-olivary projection in the rat. Anat. Embryol. (Berl). 184, 225–243.

Van Der Giessen, R.S., Koekkoek, S.K., van Dorp, S., De Gruijl, J.R., Cupido, A., Khosrovani, S., Dortland, B., Wellershaus, K., Degen, J., Deuchars, J., et al. (2008). Role of Olivary Electrical Coupling in Cerebellar Motor Learning. Neuron 58, 599–612.

Garrigues, P. J. and Ghaoui L. E. (2009). An homotopy algorithm for the Lasso with online observations. Advances in neural information processing systems, NIPS 2009.

Good J-M., Mahoney M., Miyazaki T., Tanaka K. F., Sakimura K., Watanabe M., Kitamura K. and Kano M. (2017). Maturation of Cerebellar Purkinje Cell Population Activity during Postnatal Refinement of Climbing Fiber Network. Cell Reports, 21(8):2066–2073.

Grueber, C.E., Nakagawa, S., Laws, R.J., and Jamieson, I.G. (2011). Multimodel inference in ecology and evolution: Challenges and solutions. J. Evol. Biol. 24, 699–711.

Hansel, C., Linden, D.J., and D’Angelo, E. (2001). Beyond parallel fiber LTD: the diversity of synaptic and non-synaptic plasticity in the cerebellum. Nat. Neurosci. 4, 467–475.

Herzfeld, D. J., Kojima, Y., Soetedjo, R. and Shadmehr, R. (2018). Encoding of error and learning to correct that error by the Purkinje cells of the cerebellum. Nat. Neurosci., 21: 736–743.

Hirata, Y., and Aihara, K. (2009). Representing spike trains using constant sampling intervals. J. Neurosci. Methods 183, 277–286.

Hirata, Y., Horai, S., and Aihara, K. (2008). Reproduction of distance matrices and original time series from recurrence plots and their applications. Eur. Phys. J. Spec. Top. 164, 13–22.

Hoang, H., Yamashita, O., Tokuda, I.T., Sato, M.-A., Kawato, M., and Toyama, K. (2015). Segmental Bayesian estimation of gap-junctional and inhibitory conductance of inferior olive neurons from spike trains with complicated dynamics. Front. Comput. Neurosci. 9, 1–14.

Hoge, G.J., Davidson, K.G., Yasumura, T., Castillo, P.E., Rash, J.E., and Pereda, A.E. (2011). The extent and strength of electrical coupling between inferior olivary neurons is heterogeneous. J. Neurophysiol. 105, 1089–1101.

Hoogland, T.M., De Gruijl, J.R., Witter, L., Canto, C.B., De Zeeuw, C.I. (2015). Role of synchronous activation of cerebellar Purkinje cell ensembles in multi-joint movement control. Curr Biol. 25:1157–65.

Ito, M. (2001). Cerebellar long-term depression: characterization, signal transduction, and functional roles. Physiol. Rev. 81, 1143–1195.

Kantz, H. (1994). A robust method to estimate the maximal Lyapunov exponent of a time series. Phys. Lett. A 185, 77–87.

Kaplan, J.L., and Yorke, J.A. (1970). Chaotic behavior of multidimensional difference equations. In H. O. Walter & H. Peitgen (Eds.). In Lecture Notes in Mathematics: Vol. 730. Functional Differential Equations and Approximations of Fixed Points, pp. 204–207.

Katori, Y., Lang, E.J., Onizuka, M., Kawato, M., and Aihara, K. (2010). Quantitative Modeling of Spatio-Temporal Dynamics of Inferior Olive Neurons With a Simple Conductance-Based Model. Int. J. Bifurcat. Chaos 20, 583–603.

Kawato, M., and Gomi, H. (1992). The cerebellum and VOR/OKR learning models. Trends Neurosci. 15, 445–453.

Kawato, M., Kuroda, S., and Schweighofer, N. (2011). Cerebellar supervised learning revisited: Biophysical modeling and degrees-of-freedom control. Curr. Opin. Neurobiol. 21, 791–800.

Keating, J.G., and Thach, W.T. (1995). Nonclock behavior of inferior olive neurons: interspike interval of Purkinje cell complex spike discharge in the awake behaving monkey is random. J. Neurophysiol. 73, 1329–1340.

Kitazawa, S., Kimura, T., and Yin, P.B. (1998). Cerebellar complex spikes encode both destinations and errors in arm movements. Nature 392, 494–497.

Kobayashi, Y., Kawano, K., Takemura, A., Inoue, Y., Kitama, T., Gomi, H., and Kawato, M. (1998). Temporal firing patterns of Purkinje cells in the cerebellar ventral paraflocculus during ocular following responses in monkeys II. Complex spikes. J. Neurophysiol. 80, 832–848.

Kuroda, S., Schweighofer, N., and Kawato, M. (2001). Exploration of signal transduction pathways in cerebellar long-term depression by kinetic simulation. J. Neurosci. 21, 5693–5702.

Lang, E.J. (2002). GABAergic and glutamatergic modulation of spontaneous and motor-cortex-evoked complex spike activity. J. Neurophysiol. 87, 1993–2008.

Lang E.J., Apps R., Bengtsson F., Cerminara N.L., De Zeeuw C.I., Ebner T.J., Heck D.H., Jaeger D., Jörntell H., Kawato M., Otis T.S., Ozyildirim O., Popa L.S., Reeves A.M.B., Schweighofer N., Sugihara I., Xiao J. (2016). The roles of the olivocerebellar pathway in motor learning and motor control, The Cerebellum, 16(1): 230–252.

Lang, E. J., and Blenkinsop, T. A. (2011). Control of cerebellar nuclear cells: a direct role for complex spikes? Cerebellum 10: 694–701.

Lang, E.J., Sugihara, I., and Llinás, R. (1996). GABAergic modulation of complex spike activity by the cerebellar nucleoolivary pathway in rat. J. Neurophysiol. 76, 255–275.

LeCun Y, Bengio Y, Hinton G (2015) Deep learning. Nature, 521:436–444.

Lee, K. H., Mathews, P. J., Reeves, A. M., Choe, K. Y., Jami, S. A., Serrano, R. E. and Otis, T. S. (2015). Circuit mechanisms underlying motor memory formation in the cerebellum. Neuron 86(2): 529–40

Lefler, Y., Yarom, Y., and Uusisaari, M. (2014). Cerebellar inhibitory input to the inferior olive decreases electrical coupling and blocks subthreshold oscillations. Neuron 81, 1389–1400.

Letellier, C. (2006). Estimating the Shannon entropy: Recurrence plots versus symbolic dynamics. Phys. Rev. Lett. 96, 254102.

Llinás R. The noncontinuous nature of movement execution. In: Humphrey DR, Freund H-J, editors. Motor control: concepts and issues. New York: Wiley; 1991. p. 223–42.

Llinás, R., Baker R., and Sotelo C. (1974). Electrotonic coupling between neurons in cat inferior olive. J. Neurophysiol. 37(3), 560–571.

Llinás, R., and Yarom, Y. (1981). Electrophysiology of mammalian inferior olivary neurones in vitro. Different types of voltage-dependentionic conductances. J. Physiol. 315, 549–567.

Long M.A., Deans M.R., Paul D.L., Connors B.W. (2002) Rhythmicity without synchrony in the electrically uncoupled inferior olive. J Neurosci. 22(24):10898–905.

Makarenko, V., and Llinas, R. (1998). Experimentally determined chaotic phase synchronization in a neuronal system. Proc. Natl. Acad. Sci. United States Am. Sci. 95, 15747–15752.

Marr, D. (1969). A theory of cerebellar cortex. J Physiol. 202(2): 437–470.

Marr, D. (1982). Vision: A Computational Investigation into the Human Representation and Processing of Visual Information. Henry Holt, New York, NY, USA.

Marshall S.P., van der Giessen R.S., de Zeeuw C.I. and Lang E.J. (2007). Altered olivocerebellar activity patterns in the connexin36 knockout mouse. Cerebellum. 6(4):287–99.

Marshall, S. P. and Lang E. J. (2009). Local changes in the excitability of the cerebellar cortex produce spatially restricted changes in complex spike synchrony. J Neurosci 29(45), 14352–14362.

Marwan, N., Carmen Romano, M., Thiel, M., and Kurths, J. (2007). Recurrence plots for the analysis of complex systems. Phys. Rep. 438, 237–329.

Masuda, N., and Aihara, K. (2002). Spatiotemporal spike encoding of a continuous external signal. Neural Comput. 14, 1599–1628.

Mazzucato L, Fontanini A and La Camera G (2016) Stimuli Reduce the Dimensionality of Cortical Activity. Front. Syst. Neurosci. 10:11. doi:10.3389/fnsys.2016.00011

Mnih V, Kavukcuoglu K, Silver D, Rusu A a, Veness J, Bellemare MG, Graves A, Riedmiller M, Fidjeland AK, Ostrovski G, et al. (2015). Human-level control through deep reinforcement learning. Nature, 518:529–533.

Najac, M., and Raman I. M. (2015). Integration of Purkinje Cell Inhibition by Cerebellar Nucleo-Olivary Neurons. J. Neurosci. 35(2), 544–549.

Nelson, B. J. and Mugnaini, E. (1989). Origins of GABAergic inputs to the inferior olive. In Strata, P. (ed.), The Olivocerebellur Sysrem in Motor Control. Springer-Verlag. Berlin. Exp. Bruin Res., Suppl. 17, 86–107.

Nobukawa, S., and Nishimura, H. (2016). Chaotic resonance in coupled inferior olive neurons with the Llinás approach neuron model. Neural Comput. 28, 2505–2532.

Onizuka, M., Hoang, H., Kawato, M., Tokuda, I.T., Schweighofer, N., Katori, Y., Aihara, K., Lang, E.J., and Toyama, K. (2013). Solution to the inverse problem of estimating gap-junctional and inhibitory conductance in inferior olive neurons from spike trains by network model simulation. Neural Networks 47, 51–63.

Pillow, J. W., Shlens, J., Paninski, L., Sher, A., Litke, A. M., Chichilnisky, E. J. and Simoncelli, E. P. (2008). Spatio-temporal correlations and visual signaling in a complete neuronal population. Nature 454: 995–999.

Rigotti M., Barak O., Warden M.R., Wang X.J., Daw N.D, Miller E.K. & Fusi S. (2013) The importance of mixed selectivity in complex cognitive tasks. Nature. 497:585–590.

Rosenstein, M.T., Collins, J.J., and De Luca, C.J. (1993). A practical method for calculating largest Lyapunov exponents from small data sets. Phys. D 65, 117–134.

Sano, M., and Sawada, Y. (1985). Measurement of the lyapunov spectrum from a chaotic time series. Phys. Rev. Lett. 55, 1082–1085.

Schaal, S., Atkeson, C.G. and Vijayakumar, S. (2002). Scalable Techniques from Nonparametric Statistics for Real Time Robot Learning. Applied Intelligence. 17: 49–60.

Schweighofer N. (1998). A model of activity-dependent formation of cerebellar microzones. Biological Cybernetics, 79 (2): 97–107.

Schweighofer, N., Spoelstra, J., Arbib, M.A., and Kawato, M. (1998). Role of the cerebellum in reaching movements in humans. II. A neural model of the intermediate cerebellum. Eur. J. Neurosci. 10, 95–105.

Schweighofer, N., Doya, K., and Kawato, M. (1999). Electrophysiological properties of inferior olive neurons: A compartmental model. J. Neurophysiol. 82, 804–817.

Schweighofer, N., Doya, K., Fukai, H., Chiron, J.V., Furukawa, T., and Kawato, M. (2004). Chaos may enhance information transmission in the inferior olive. Proc. Natl. Acad. Sci. 101, 4655–4660.

Schweighofer, N., Lang, E.J., and Kawato, M. (2013). Role of the olivo-cerebellar complex in motor learning and control. Front. Neural Circuits 7, 94.

Shaikh, A. G., Wong, A. L., Optican, L. M. and Zee, D. S. (2017) Impaired Motor Learning in a Disorder of the Inferior Olive: Is the Cerebellum Confused? Cerebellum 16: 158–167.

Shew W.L., Yang H., Yu S., Roy R. and Plenz D. (2011) Information Capacity and Transmission Are Maximized in Balanced Cortical Networks with Neuronal Avalanches. J. Neurosci. 31(1) 55–63.

Shinomoto, S., Miura, K., and Koyama, S. (2005). A measure of local variation of inter-spike intervals. Bio Systems, 79(1-3): 67–72.

Silver D, Huang A, Maddison CJ, Guez A, Sifre L, van den Driessche G, Schrittwieser J, Antonoglou I, Panneershelvam V, Lanctot M, et al. (2016). Mastering the game of Go with deep neural networks and tree search. Nature, 529:484–489.

Sotelo, C., Llinás R., and Baker R. (1974). Structural study of inferior olivary nucleus of the cat: morphological correlates of electrotonic coupling. J. Neurophysiol. 37(3), 541–559.

Sugihara, I., Fujita, H., Na, J., Quy, P.N., Li, B.Y. and Ikeda, D. (2009). Projection of reconstructed single Purkinje cell axons in relation to the cortical and nuclear aldolase C compartments of the rat cerebellum. J Comp Neurol. 512(2):282–304.

Sugihara, I. and Shinoda, Y. (2004). Molecular, topographic, and functional organization of the cerebellar cortex: a study with combined aldolase C and olivocerebellar labeling. J Neurosci. 24(40): 8771–85.

Sugihara, I. and Shinoda, Y. (2007). Molecular, Topographic, and Functional Organization of the Cerebellar Nuclei: Analysis by Three-Dimensional Mapping of the Olivonuclear Projection and Aldolase C Labeling. J Neurosci. 27(36): 9696–9710.

Tang, T., Suh C. Y., Blenkinsop T. A., and Lang E. J. (2016). Synchrony is Key: Complex Spike Inhibition of the Deep Cerebellar Nuclei. Cerebellum 15(1), 10–13.

Tokuda, I.T., Han, C.E., Aihara, K., Kawato, M., and Schweighofer, N. (2010). The role of chaotic resonance in cerebellar learning. Neural Networks 23, 836–842.

Tokuda, I.T., Hoang, H., Schweighofer, N., and Kawato, M. (2013). Adaptive coupling of inferior olive neurons in cerebellar learning. Neural Networks 47, 42–50.

Tokuda, I.T., Hoang, H., and Kawato, M. (2017). New insights into olivo-cerebellar circuits for learning from a small training sample, Curr. Opin. Neurobiol. 46:58–67.

Tseng, Y.-W., Diedrichsen, J., Krakauer, J.W., Shadmehr, R., and Bastian, A.J. (2007). Sensory prediction errors drive cerebellum-dependent adaptation of reaching. J. Neurophysiol. 98, 54–62.

Turecek, J., Yuen, G.S., Han, V.Z., Zeng, X.H., Bayer, K.U., and Welsh, J.P. (2014). NMDA receptor activation strengthens weak electrical coupling in mammalian brain. Neuron 81, 1375–1388.

Victor, J.D., and Purpura, K.P. (1997). Metric-space analysis of spike trains: theory, algorithms and application. Netw. Comput. Neural Syst. 8, 127–164.

Vinueza Veloz, M.F., Zhou, K., Bosman, L.W.J., Potters, J.W., Negrello, M., Seepers, R.M., Strydis, C., Koekkoek, S.K.E., and De Zeeuw, C.I. (2015). Cerebellar control of gait and interlimb coordination. Brain Struct. Funct. 220, 3513–3536.

Watanabe S (2009). Algebraic Geometry and Statistical Learning Theory. Cambridge University Press.

Welsh, J.P., Lang, E.J., Sugihara, I., Llinás, R. (1995). Dynamic organization of motor control within the olivocerebellar system. Nature 374:453–7.

Wilkinson, G.N., and Rogers, C.E. (1973). Symbolic description of factorial models for analysis of variance. Appl. Stat. 22, 392–399.

Witter, L., Canto, C.B., Hoogland, T.M., de Gruijl, J.R., and De Zeeuw, C.I. (2013). Strength and timing of motor responses mediated by rebound firing in the cerebellar nuclei after Purkinje cell activation. Front. Neural Circuits 7: 133.

Wolf, A., Swift, J.B., Swinney, H.L., and Vastano, J.A. (1985). Determining Lyapunov exponents from a time series. Phys. D Nonlinear Phenom. 16, 285–317.

Yamazaki K (2014). Asymptotic accuracy of distribution-based estimation of latent variables. J Mach Learn Res. 15:3721–3742.

De Zeeuw, C.I., and Ten Brinke, M.M. (2015). Motor learning and the cerebellum. Cold Spring Harb. Perspect. Biol. 7.

De Zeeuw, C.I., Holstege, J.C., Ruigrok, T.J.H., and Voogd, J. (1989). Ultrastructural study of the GABAergic, cerebellar, and mesodiencephalic innervation of the cat medial accessory olive: Anterograde tracing combined with immunocytochemistry. J. Comp. Neurol. 284, 12–35.

De Zeeuw, C.I., Hertzberg, E.L., and Mugnaini, E. (1995). The dendritic lamellar body: a new neuronal organelle putatively associated with dendrodendritic gap junctions. J. Neurosci. 15,1587–1604.

De Zeeuw, C.I., Simpson, J.I., Hoogenraad, C.C., Galjart, N., Koekkoek, S.K.E., and Ruigrok, T.J.H. (1998). Microcircuitry and function of the inferior olive. Trends Neurosci. 21, 391–400.

